# Aβ*56 is a stable oligomer that correlates with age-related memory loss in Tg2576 mice

**DOI:** 10.1101/2023.03.20.533414

**Authors:** Peng Liu, Ian P. Lapcinski, Samantha L. Shapiro, Lisa J. Kemper, Karen H. Ashe

## Abstract

Amyloid-β (Aβ) oligomers consist of fibrillar and non-fibrillar soluble assemblies of the Aβ peptide. Tg2576 human amyloid precursor protein (APP)-expressing transgenic mice modeling Alzheimer’s disease produce Aβ*56, a non-fibrillar Aβ assembly that has been shown by several groups to relate more closely to memory deficits than plaques. Previous studies did not decipher specific forms of Aβ present in Aβ*56. Here, we confirm and extend the biochemical characterization of Aβ*56. We used anti-Aβ(1-x), anti-Aβ(x-40), and A11 anti-oligomer antibodies in conjunction with western blotting, immunoaffinity purification, and size-exclusion chromatography to probe aqueous brain extracts from Tg2576 mice of different ages. We found that Aβ*56 is a ∼56-kDa, SDS-stable, A11-reactive, non-plaque-related, water-soluble, brain-derived oligomer containing canonical Aβ(1-40) that correlates with age-related memory loss. The unusual stability of this high molecular-weight oligomer renders it an attractive candidate for studying relationships between molecular structure and effects on brain function.

## Introduction

Aβ*56 was the first brain-derived Aβ oligomer shown to impair memory in mice and rats^1^. Aβ*56 forms before plaques appear, and is found both inside dense-core plaques and dispersed outside in the brain parenchyma^2^. The discovery of Aβ*56 stimulated interest in Aβ oligomers as mediators of cognitive deficits in Alzheimer’s disease (AD), and spurred the development of new, oligomer-specific, therapeutic antibodies. Aβ*56 is but one of many variants of brain-derived Aβ oligomers. Other brain-derived, oligomeric variants include dimers and trimers^3^, prefibrillar oligomers that bind A11 antibodies^4^, amylospheroids^5^, globulomers^6^, fibrillar oligomers that bind OC antibodies^7^, protofibrillar oligomers that bind mAb158/lecanumab antibodies^8^, intraneuronal oligomers that bind NU-1 anti-ADDL antibodies^9^, annular protofibrils^10^, amyloid pores^11^, oligomers that bind crenezumab^12^, oligomers that bind aducanumab^13^, oligomers that bind ACU3B3/ACU193 anti-ADDL antibodies^14^, oligomers that bind α-sheets^15^, and oligomers that bind JD1 antibodies^16^, whose structure, spatial distribution, temporal expression, biogenesis, and effects on brain function are topics of active study with important therapeutic implications.

Aβ*56 correlates with aging and impaired memory in mice, dog, and humans. Billings *et al* found that repeated behavioral training decreases Aβ*56 levels and amyloid plaque loads and improves memory function, reflecting a possible connection between brain activity and Aβ aggregation^17^. Pop *et al* showed a positive correlation between age and Aβ*56 in aqueous extracts of temporal cortex from beagles^18^. Yoo *et al* demonstrated an association between Aβ*56 and age-related apoptosis in the ectoturbinate olfactory epithelium of Tg2576^19^ mice^20^. Building upon these findings, Yoo *et al* investigated nasal fluid in living, elderly human participants and found a ∼2.9-fold increase in Aβ*56 in individuals diagnosed with AD compared to controls^21^. In addition, over three years, subjects with Aβ*56 levels above the median value deteriorated while those with levels below the median remained stable^21^.

Studies in different lines of APP transgenic mice indicate that memory deficits relate more closely to Aβ*56 than to dense-core amyloid plaques or Aβ dimers, the smallest oligomeric constituent of dense-core plaques identified under denaturing sodium dodecyl sulfate/polyacrylamide gel electrophoresis (SDS-PAGE) conditions^22^. Cheng *et al* demonstrated that spatial reference memory in J20^23^ mice expressing moderate levels of APP-V717F negatively correlates with levels of Aβ*56 but not dense-core plaque loads^24^. Arc6^25^ mice, which express low levels of APP-V717F with the E693G Arctic mutation promoting fibril formation leading to plaque loads that are higher than those in J20 mice, lack Aβ*56 and, accordingly, exhibit normal spatial reference memory^24^. Meilandt *et al* showed that neprilysin overexpression in J20 mice reduces amyloid plaques but not Aβ*56, and does not improve spatial reference memory, elevated plus-maze performance, or premature mortality^26^. Liu *et al* showed that grape-seed polyphenolic extracts in drinking water, previously shown to attenuate cognitive impairment in Tg2576 mice^27^, reduces levels of Aβ*56 by 48% but did not change levels of Aβ dimers^28^. Castillo-Carran *et al* reported improved performance in contextual fear conditioning and novel object recognition in Tg2576 mice two weeks after a single intravenous injection of tau-oligomer monoclonal antibodies is accompanied by ∼60% reduction in Aβ*56 level and ∼180% increase in ThioS amyloid plaque load^29^, suggesting that tau oligomers promote off-pathway and depress on-pathway fibril aggregation of Aβ. Liu *et al* furnished a rationale for the dissociation between dense-core amyloid plaques and impaired cognition by demonstrating that Aβ dimers, which are confined to dense-core plaques in Tg2576 and rTg9191^30^ mice, do not impair memory function unless they are extracted from the plaques and dispersed in the brain^2, 30^.

Here, we confirm and extend the characterization of Aβ*56, and show that it is a ∼56-kDa, A11-reactive, SDS-stable, non-plaque-related, water-soluble, brain-derived oligomer that contains canonical Aβ(1-40) and correlates with age-related memory loss in Tg2576 mice.

## Results

### Age-related appearance of a ∼56-kDa, SDS-stable, water-soluble, Aβ-containing entity in Tg2576 mice

We screened proteins from aqueous brain extracts immunoprecipitated using anti-Aβ(x-40) antibodies by fractionating them using SDS-PAGE, and probing them on western blots using 6E10 antibodies recognizing Aβ(3-8) (**Fig. 1A**). We screened both plaque-free (≤12 months of age)^31^ and plaque-bearing (≥13 months of age)^31^ Tg2576 mice, and found a ∼56-kDa, 6E10-reactive entity in a subset of Tg2576 mice, but not in non-transgenic littermates or rTg9191 mice (**Fig. 1B; Table S1**). We detected no other specific, 6E10-reactive entities except the ∼56-kDa entity and ∼4.5-kDa monomeric Aβ. The ∼56-kDa, 6E10-reactive entity was present in 0% of 2-5 month, 30% of 6-11 month, 65% of 12-16 month, and 100% of 21-24 month mice (**Fig. 1C; Table S1**). These findings are temporally consistent with the previously reported onset and progression of memory loss in Tg2576 mice (**Fig. 1D**), whose memory function begins to deteriorate at 6 months and gradually worsens with age^32^. To designate mice with impaired memory function, we applied a cut-off of 35% mean probe score (MPS), which generates nominal values from a continuous, distributed variable; this likely explains why some non-transgenic and 2-5 month mice appear impaired, and not all of the 21-24 month mice appear impaired. We concluded that a water-soluble, SDS-stable, Aβ-containing entity that migrates at ∼56 kDa is present in Tg2576 mice, and that its initial appearance and rising prevalence correspond to the onset of memory loss and progressive deterioration in memory function characteristic of Tg2576 mice.

**Figure 1.**
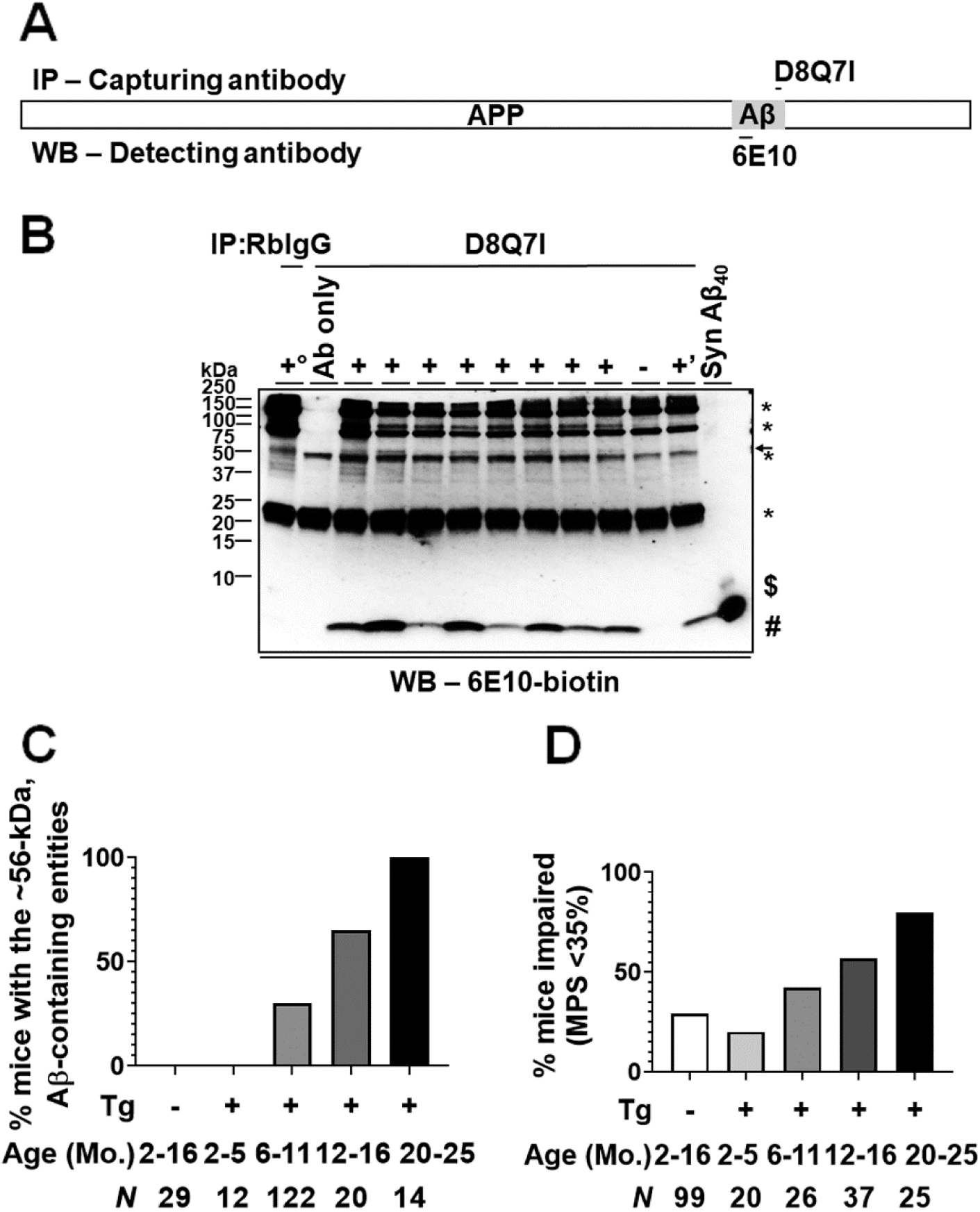
Age-related presence of ∼56-kDa, SDS-stable, water-soluble, Aβ-containing entities in Tg2576 mice. **A**) A schematic illustration of epitopes of capturing and detecting antibodies used in immunoprecipitation (IP)/Western blotting (WB) for panel B. Capturing antibody: rabbit monoclonal D8Q7I that is directed against the C-terminus of Aβ(x-40), detecting antibody: biotinylated mouse monoclonal 6E10 that is directed against Aβ(3-8). **B**) A representative IP/WB analysis showing that D8Q7I-precipitated entities electrophoresed at ∼56 kDa (arrow) are detected by biotinylated 6E10 (6E10-biotin) in a subset of Tg2576 (+) but not in a non-transgenic littermate of Tg2576 mice (-), in an rTg9191 mouse (+’), or when no Tg2576 brain extracts were used (Ab only). No ∼56-kDa entities are detected when generic rabbit immunoglobulin G (RbIgG) was used to precipitate brain extracts (+°, combined brain extracts of the eight Tg2576 mice, each contributing 12.5% of the total proteins). Ab, capturing antibody D8Q7I. APP, amyloid precursor protein. Aβ, amyloid-β. Syn Aβ_40_, synthetic Aβ(1-40) (5 ng), serving as a positive control for WB. #, monomeric Aβ; $, dimeric Syn Aβ_40_; and *, non-specifically detected entities. kDa, kilo-Daltons. **C**) The percentage of Tg2576 mice expressing ∼56-kDa entities increases with age. Non-transgenic littermates (-) and 2-5 month Tg2576 mice (+) do not express the ∼56-kDa entity. **D**) The percentage of Tg2576 mice exhibiting impaired spatial reference memory increases with age. The percentage of impaired 2-5 month Tg2576 mice (+) is similar to that of 2-16 month, non-transgenic littermates (-), representing baseline-deficits. This analysis was performed using previously described data acquired using a water maze.^32^ Impaired spatial reference memory is defined here as a mean probe score (MPS) <35%.^32^ The sample size for each age period is shown. Mo., months. Tg, transgene expressing human amyloid precursor protein.

### Canonical Aβ(1-40) is present in the ∼56-kDa, SDS-stable, water-soluble, Aβ-containing entities

We captured proteins in aqueous brain extracts by immunoprecipitation (IP) with D8Q7I antibodies recognizing the C-terminus of Aβ(x-40), fractionated captured proteins by SDS-PAGE, transferred proteins to nitrocellulose membranes, and probed them using biotinylated 82E1 antibodies recognizing the N-terminus of Aβ(1-x) (**Fig. 2A**). Immunoprecipitated proteins probed with 82E1 antibodies revealed a ∼56-kDa entity in 8-12 month, plaque-free, Tg2576 mice, but not in non-transgenic littermates (**Fig. 2B**). We detected monomeric Aβ in all Tg2576 mice, and an entity migrating at ∼40 kDa in some mice. Using the same methodology, we also found ∼56-kDa entities containing canonical Aβ(1-40) in 14-21 month, plaque-bearing Tg2576 mice (**Fig. S1**). We concluded that the ∼56-kDa, SDS-stable, water-soluble entities in Tg2576 mice contain canonical Aβ(1-40).

**Figure 2.**
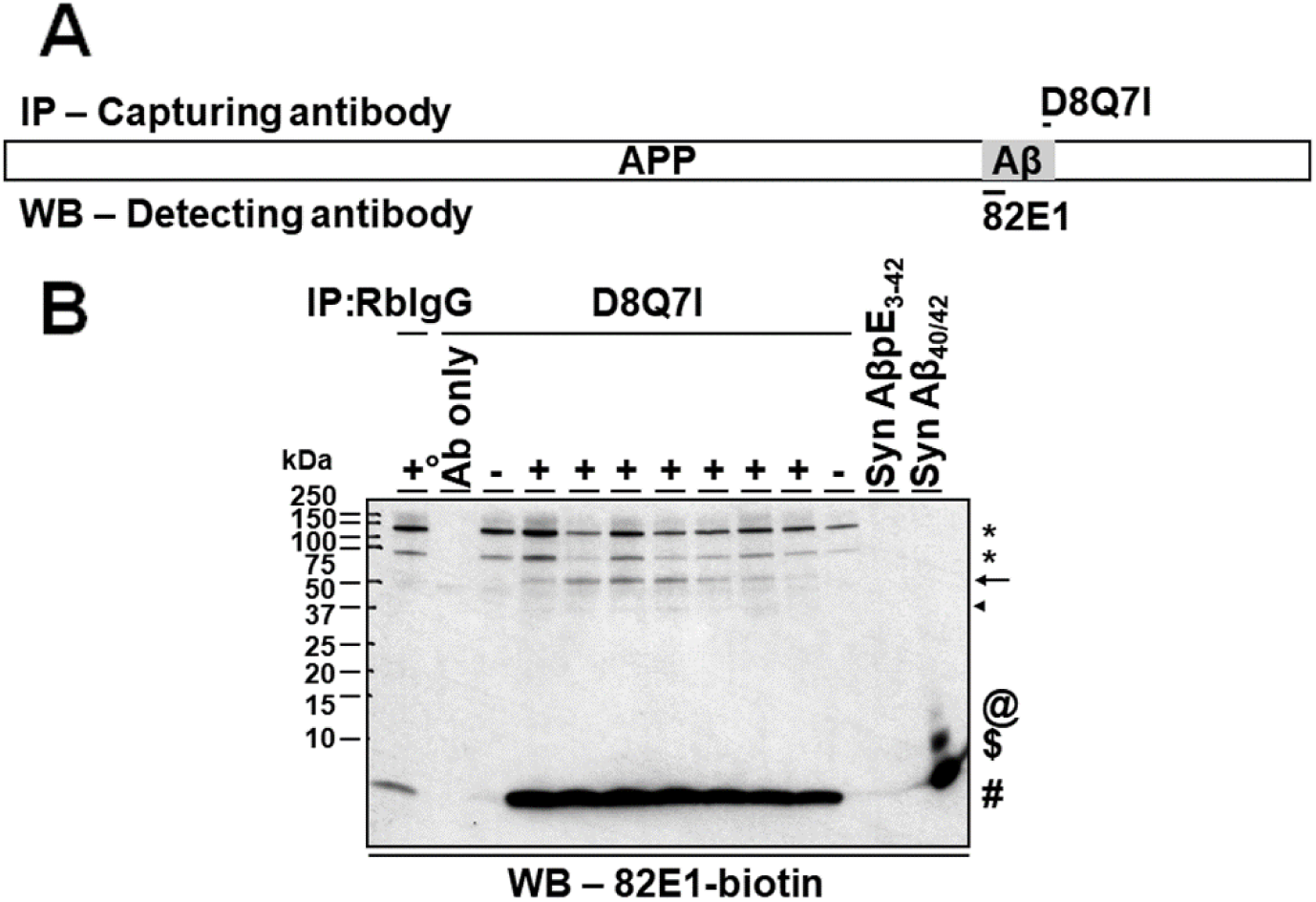
Canonical Aβ(1-40) is present in the ∼56-kDa, SDS-stable, water-soluble, Aβ-containing entities in Tg2576 mice. **A**) A schematic illustration of epitopes of capturing and detecting antibodies used in immunoprecipitation (IP)/western blotting (WB) for panel B. Capturing antibody: rabbit monoclonal D8Q7I, detecting antibody: biotinylated mouse monoclonal 82E1 that is directed against the N-terminus of Aβ(1-x). **B**) A representative IP/WB analysis showing that D8Q7I-precipitated entities electrophoresed at ∼56 kDa (arrow) are detected by biotinylated 82E1 (82E1-biotin) in 8-12 month Tg2576 mice (+) but not in age-matched, non-transgenic littermates (-). In addition, ∼40-kDa entities (arrowhead) in some Tg2576 mice are also detected. No ∼56-kDa, D8Q7I-precipitated entities are detected when no Tg2576 brain extracts were used (Ab only). No ∼56-kDa entities are detected in Tg2576 mice when generic rabbit immunoglobulin G (RbIgG) was used to precipitate brain extracts (+°, combined brain extracts of the seven Tg2576 mice, each contributing 14.3% of the total proteins). Ab, capturing antibody D8Q7I. APP, amyloid precursor protein. Aβ, amyloid-β. Syn AβpE3-42, synthetic Aβ_42_ with the 3^rd^ amino acid being a pyroglutamate (30 ng), serving as a negative control for WB; and Syn Aβ_40/42_, synthetic Aβ(1-40) (5 ng) and Aβ(1-42) (5 ng), serving as a positive control for WB. #, monomeric Aβ; $, dimeric Syn Aβ_40/42_; @, trimeric Syn Aβ_40/42_; and *, non-specifically detected entities. kDa, kilo-Daltons.

Next, we reversed the order of the antibodies used for IP and western blotting. Proteins immunoprecipitated with 82E1 antibodies that were probed on western blots with biotinylated D8Q7I antibodies did not reveal ∼56-kDa assemblies (**Fig. S2**), suggesting that the free N-terminus is inaccessible under non-denaturing conditions.

Some forms of Aβ aggregate to form amyloid fibrils more readily than Aβ(1-40), notably Aβ(x-42)^33^. We sought, therefore, to determine whether the ∼56-kDa entity contains Aβ(x-42). We captured proteins in brain extracts of the same plaque-free Tg2576 mice studied above using D3E10 antibodies recognizing Aβ(x-42), and analyzed the captured proteins in western blots probed with biotinylated 6E10 antibodies. We detected no ∼56-kDa bands in these specimens (**Fig. S3**). We confirmed these results using Aβ(x-40) and Aβ(x-42) antibodies from another source (**Fig. S4**). These studies indicate that, within the limits of our assay, the ∼56-kDa, Aβ-containing entity in Tg2576 mice contains no detectable Aβ(x-42).

### The ∼56-kDa, Aβ(1-40)-containing entities are not artificially formed by exposure to SDS

Although our results indicate that the ∼56-kDa entities contain canonical Aβ(1-40), these data do not exclude the possibility that they are artificially generated from monomers exposed to SDS. It is also possible that they are not oligomers, but rather complexes of Aβ(1-40) monomers covalently bound together or to other macromolecules.

To investigate the possibility that the ∼56-kDa, Aβ(1-40)-containing entity may be artificially generated from monomers exposed to SDS, we used non-denaturing size-exclusion chromatography (SEC) to fractionate aqueous brain extracts from plaque-free Tg2576 mice, and examined every other eluted fractions by SDS-PAGE and western blotting with 82E1 and D8Q7I antibodies. Numerous non-specific bands represent non-specific binding to Neutravidin (**Fig. S5**).

We found most of the 82E1-reactive, ∼56-kDa entities in fractions corresponding to ∼46-120 kDa (Fractions 59-67) with a peak at ∼55 kDa, but not in the fractions containing monomeric Aβ, which appeared in fractions corresponding to ∼160 to >2,000 kDa and ∼10-24 kDa (Fractions 37-57 and 75-81, respectively) (**Figs. 3A and 3B**). The early peak of monomers reflects the presence of large aggregates containing weakly associating monomers, while the late peak contains small, SDS-sensitive assemblies, findings that are consistent with a previously published result ^34^. A minor portion of the ∼56-kDa entities were found in fractions corresponding to ∼200-540 kDa (Fractions 47-55) with a peak at ∼290 kDa. The presence of the ∼56-kDa entity in fractions corresponding to higher molecular masses suggests that under non-denaturing conditions, some ∼56-kDa entities bind weakly together. A specific, ∼14-kDa band appeared in fractions corresponding to ∼24-40 kDa (Fractions 69-75) with a peak at ∼29 kDa.

**Figure 3.**
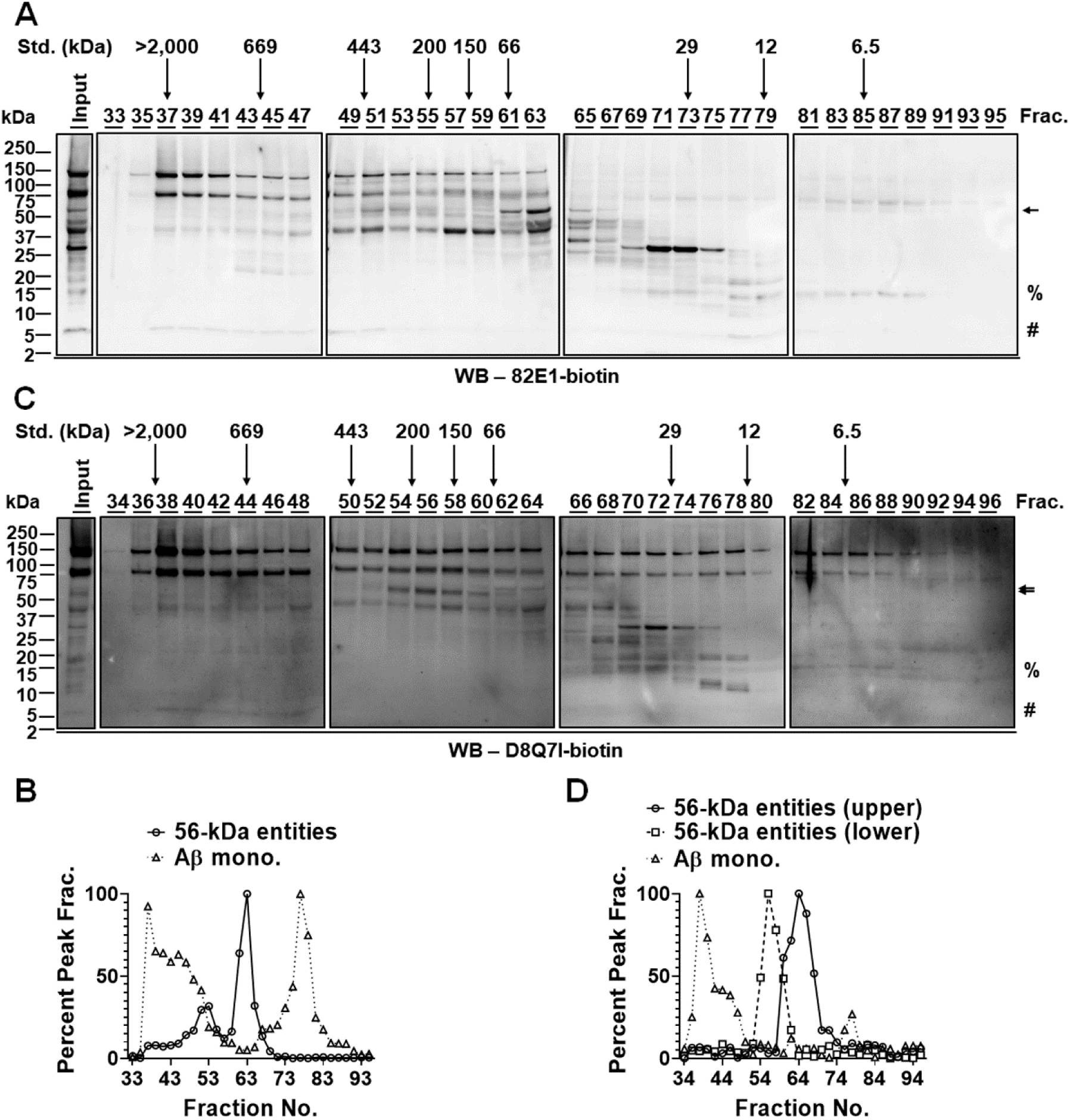
The ∼56-kDa, Aβ(1-40)-containing entities are not artificially formed by exposure to SDS. **A**) Representative western Blotting (WB) analyses showing the detection of the ∼56-kDa, Aβ-containing entities (arrow, fractions (Frac.) 47-67), ∼14-kDa entities (%, Frac. 69-75), and monomeric Aβ (#, Frac. 37-55 and 75-81) in odd-numbered size-exclusion chromatography (SEC) fractions (Frac. 33-95) of 10-11 month Tg2576 brain extracts using biotinylated 82E1 (82E1-biotin). The four WB images (i.e., Frac. 33-47, 49 to 63, 65-79, and 81-95) shown were obtained using the same exposure time (20 sec), and the WB band patterns and intensities of the input material (5% (w/v) of the brain extracts used for SEC) within each of the four WB images are comparable. A WB of the input is shown in the far left panel. The elution profile of biomolecule standards (Std.) of varied sizes is shown above the WB images. kDa, kilo-Daltons. **B**) Quantitative analyses of levels of the ∼56-kDa, 82E1-reactive entities and monomeric Aβ in SEC fractions. The level of the ∼56-kDa entities in each fraction is normalized to its highest level in fraction 63, and of monomeric Aβ (Aβ mono.) in each fraction is normalized to its highest level in fraction 77. **C**) Representative WB analyses showing the detection of the ∼56-kDa, Aβ-containing entities (arrows, Frac. 52-72), ∼14-kDa entities (%, Frac. 68-74), and monomeric Aβ (#, Frac. 36-48 and 76-78) in even-numbered SEC fractions (Frac. 34-96) of 10-11 month Tg2576 brain extracts using biotinylated D8Q7I (D8Q7I-biotin). The four WB images (i.e., Frac. 34-48, 50-64, 66-80, and 82-96) were obtained using the same exposure time (50 sec), and the WB band patterns and intensities of the input material (5% (w/v) of the brain extracts used for SEC) within each of the four WB images are comparable. A WB of the input is shown in the far left panel. The elution profile of biomolecule standards (Std.) of varied sizes is shown above the WB images. kDa, kilo-Daltons. **D**) Quantitative analyses of levels of the ∼56-kDa, D8Q7I-reactive entities and monomeric Aβ in SEC fractions. Levels of the ∼56-kDa entities in each fraction are normalized to their highest level in fraction 56 (lower band) and fraction 64 (upper band), and the level of monomeric Aβ (Aβ mono.) in each fraction is normalized to its highest level in fraction 38.

When probed with D8Q7I antibodies, we found a doublet at ∼56 kDa (**Figs. 3C and 3D**). The upper band was present in fractions corresponding to ∼32-87 kDa (Fractions 60-72) with a peak at ∼55 kDa, while the lower band was present in fractions corresponding to ∼62-335 kDa (Fractions 52-62) with a peak at ∼180 kDa. The ∼56-kDa bands were absent in fractions containing monomeric Aβ, which appeared in fractions corresponding to ∼500 to >2,000 kDa and ∼17.5-20 kDa (Fractions 36-48 and 76-78, respectively). A specific, ∼14-kDa band appeared in fractions corresponding to ∼26-45 kDa (Fractions 68-74) with a peak at ∼28 kDa.

These results indicate that a ∼56-kDa entity that is not artificially formed from monomers or lower molecular-weight Aβ species exists, and that its mass is similar whether measured by SEC or SDS-PAGE. The ∼14-kDa band may represent a trimeric form of Aβ that weakly binds to other trimers to form a hexamer. The constituents in the D8Q7I-reactive, ∼56-kDa doublets are unclear; it is possible that they reflect heterogeneity in the N-termini of the constituent Aβ species that assemble to form hetero-oligomers.

### The ∼56-kDa, Aβ(1-40)-containing entities dissociate in denaturants

To determine whether the ∼56-kDa entity is a complex of Aβ(1-40) molecules that are covalently bound to themselves or other macromolecules, we subjected desiccated proteins in aqueous brain extracts to urea, guanidine hydrochloride (GuHCl), and 1,1,1,3,3,3-hexafluoro-2-propanol (HFIP). We resuspended the denatured proteins in Tris-buffered saline (TBS), and analyzed them by IP using D8Q7I antibodies followed by western blotting using 82E1 antibodies. Non-denatured, captured proteins (exposed to TBS only) probed with 82E1 antibodies revealed not only a ∼56-kDa band, but also bands at ∼40 kDa, ∼35 kDa, and ∼12 kDa, in addition to monomers (**Fig. 4**). All four bands disappeared after the captured proteins were denatured using urea or GuHCl, indicating that the proteins in these bands are not covalently linked (**Fig. 4**). We cannot exclude the possibility that a macromolecule is non-covalently bound to Aβ(1-40) in a manner that resists SDS. Arguing against this possibility is that the ∼56-kDa, ∼40-kDa, ∼35-kDa, and ∼12-kDa entities would require Aβ(1-40) to be complexed to a different macromolecule in each band, which seems unlikely. We concluded that the ∼56-kDa entity is an SDS-stable, water-soluble, brain-derived oligomer containing canonical Aβ(1-40), also known as Aβ*56.

**Figure 4.**
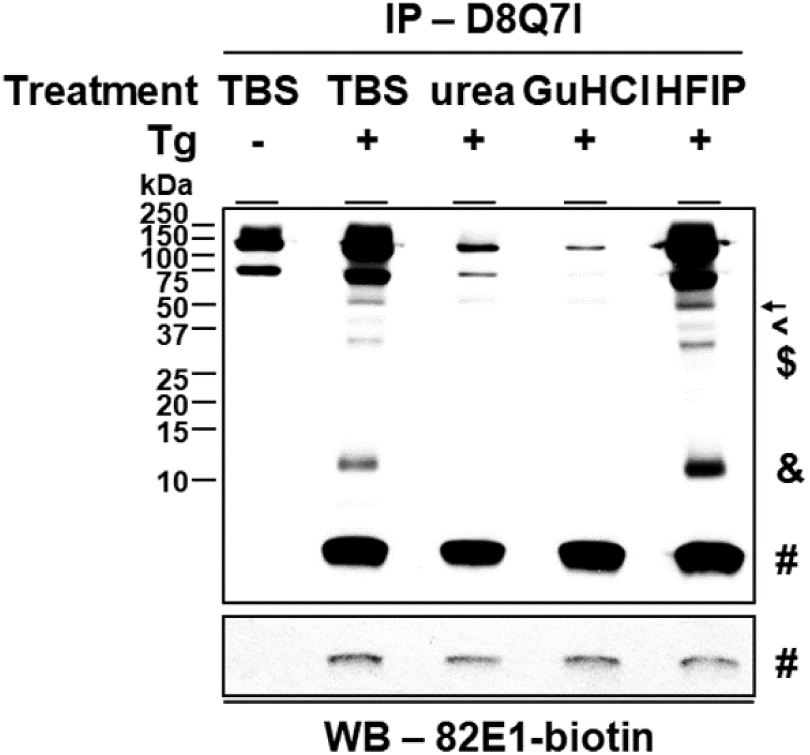
The ∼56-kDa, Aβ(1-40)-containing entities are dissociated by denaturants. A representative immunoprecipitation (IP)/western blotting (WB) analysis showing the detection of a set of 82E1-reactive Aβ(1-40)-containing entities — the ∼56-kDa (arrow), ∼40-kDa (<), ∼35-kDa ($), and ∼12-kDa (&), and monomeric Aβ (#)—in D8Q7I-precipitated brain extracts of 8-11 month Tg2576 mice (+) but not age-matched littermates (-). The detection of these entities, except for monomeric Aβ, is diminished after treating the D8Q7I-precipitated molecules with either 8M urea or 6M guanidine hydrochloride (GuHCl) but not 100% (v/v) 1,1,1,3,3,3-hexafluoro-2-propanol (HFIP), compared to the samples treated with Tris-buffered saline (TBS). Monomeric Aβ is similarly detected following these different treatments. Upper panel, light exposure; and lower panel, heavy exposure. Tg, transgene expressing human amyloid precursor protein. kDa, kilo-Daltons.

### Isolated Aβ*56 is a stable, A11-reactive, ∼56-kDa oligomer containing Aβ(1-40)

To investigate the properties of isolated Aβ*56, we immunopurified Aβ(x-40) species from aqueous brain extracts of 15-21 month Tg2576 mice using a D8Q7I-immobilized immunoaffinity matrix. We estimate that 1 mg aqueous brain protein extracts contain 1 ng Aβ*56, or 1 part per million by mass. We used 82E1 and A11 antibodies to probe immunoblots of Aβ(x-40) species, which were not detectable in the absence of these antibodies (**Fig. S6**). When probed with 82E1 antibodies, we found bands corresponding to ∼56 kDa, between 10 and 50 kDa, and monomers, indicating that these species contain canonical Aβ(1-40) (**Fig. 5A**). These data show that Aβ*56 can exist as a stable complex that resists boiling in β-mercaptoethanol and exposure to SDS.

**Figure 5.**
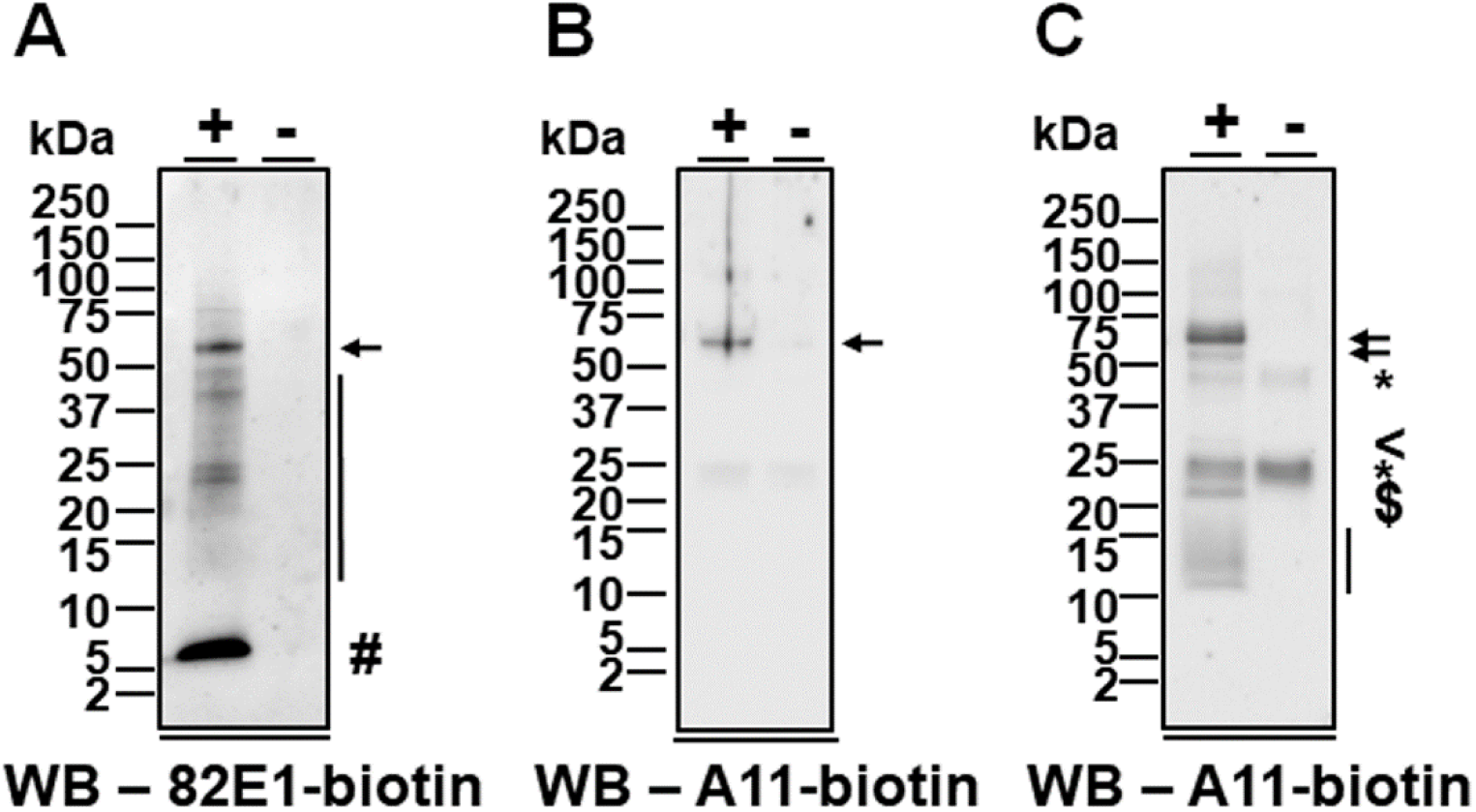
Immunopurified Aβ*56 is SDS-stable, contains canonical Aβ(1-40), and is recognized by A11, anti-oligomer antibodies. **A)** A representative western blotting (WB) analysis showing the detection of the ∼56-kDa, D8Q7I-purified, 82E1-reactive entities (arrow), monomeric Aβ (#), and multiple entities of intermediate sizes (10-50 kDa, vertical line) in the brains of 15-21 month Tg2576 mice (+) but not age-matched littermates (-). **B)** A representative WB analysis showing that under sodium dodecyl sulfate (SDS) denaturing conditions, the ∼56-kDa, D8Q7I-purified entities (arrow) but no other entities bind to the biotinylated anti-oligomer antibody A11 (A11-biotin) in the brains of 15-21 month Tg2576 mice (+) but not age-matched littermates (-). **C**) A representative WB analysis showing that under SDS semi-denaturing conditions, the D8Q7I-purified, ∼56-kDa (arrows), ∼27-kDa (<), ∼22-kDa ($), and ∼10-15-kDa (vertical line) entities are reactive to biotinylated anti-oligomer antibody A11 (A11-biotin) in the brains of 15-21 month Tg2576 mice (+) but not age-matched littermates (-). kDa, kilo-Daltons. *, non-specifically detected entities.

Next, we probed immunopurified Aβ(x-40) species using polyclonal, anti-oligomer, A11 antibodies. We previously showed that A11 does not recognize anti-parallel or in-register, parallel β-sheets, and binds entities in plaque-free J20 and Tg2576 APP transgenic mice^2^. We tested four different batches and found that A11 from two of batches detects ∼56-kDa entities (**Fig. S7**). We fractionated immunopurified Aβ(x-40) species by SDS-PAGE under denaturing (**Fig. 5B**) or semi-denaturing (**Fig. 5C**) conditions, and found a ∼56-kDa, A11-reactive band under both conditions. Under denaturing conditions (**Fig. 5B**), Aβ*56 was the sole A11-reactive species, while under semi-denaturing condition (**Fig. 5C**), there were additional species migrating at ∼27 kDa, ∼22 kDa, and ∼10-15 kDa, which likely represent less stable Aβ(1-40) assemblies.

## Discussion

In this paper, we show that Aβ*56 is a ∼56-kDa, A11-reactive, SDS-stable, water-soluble, non-plaque-related, brain-derived oligomer containing canonical Aβ(1-40) that correlates with age-related memory loss in Tg2576 mice. Aβ*56 is the only consistently present, high molecular-weight, SDS-stable, brain-derived oligomer that we were able to detect using the current experimental settings. We found that Aβ*56 can be sufficiently stable such that it does not undergo noticeable changes in conformation or size when boiled in reducing agents and exposed to SDS. To our knowledge, no other high molecular-weight Aβ oligomers exhibiting this degree of stability have been described.

The term Aβ* pays homage to Charles Weissmann’s moniker for a hypothetical form of the prion protein (PrP) that is “an infectious entity [which] could be a subspecies of PrP^Sc^ or a different modification of PrP altogether (which one might call PrP*)”^35^. Conceptually an essential component of the prion, PrP* is by definition neurotoxic and could potentially be involved in the replication of prions^36^. More recent studies indicate, however, that prion neurotoxicity and infectivity can be dissociated^37^. In keeping with PrP*, Aβ*56 appears to be neurotoxic and associates closely with impaired memory^1, 17, 18, 20, 21, 24, 26, 28, 29, 38^, but whether it is involved in the replication, aggregation, or propagation of Aβ is unclear. Aβ*56 is an A11-reactive oligomer, but has not yet been shown to behave like synthetic, A11-reactive, “prefibrillar” or α-sheet-binding Aβ oligomers in solution, which evolve to become A11-negative, fibrillar oligomers and fibrils^39, 40^. To explain the inverse relationships between insoluble Aβ and memory function that are observed only when mice are stratified by age, Westerman *et al* postulated the existence of a soluble, intermediate Aβ aggregate in Tg2576 mice that both impairs memory and facilitates the production of insoluble Aβ species and amyloid plaques^32^. Clarifying the relationship between this hypothetical Aβ intermediate and Aβ*56 will require further investigation.

While many groups have successfully studied Aβ*56 in mice^1, 17, 20, 24, 26, 28, 29^, aging dogs^18^, and humans diagnosed with AD^21, 38^, detecting Aβ*56 can be challenging because it is rare (∼1 part per million by mass). We have found that high background and non-specific binding are the greatest impediments to accurate detection. In the current studies, we have modified the original protocol developed 20 years ago to reduce non-specific signals. Using this protocol, an independent laboratory detected Aβ*56 without difficulty (**Fig. S8**). It is worth mentioning a few examples of the kinds of optimized experimental parameters that enhance the sensitivity and specificity of signals sufficiently to detect Aβ*56 reliably. We adjusted the buffers used to extract proteins, because we found that extracts containing detergents form irreversible, insoluble precipitates and may contain membrane-associated proteins. We found that tricine in gels accentuates the compactness of the Aβ*56 band; the corresponding bands in gels lacking tricine are more diffuse and less intense. The use of monoclonal antibodies that have been purified using protein A to capture proteins obscures Aβ*56 due to a non-specific band at ∼50 kDa caused by protein A contaminants in the precipitated proteins^41^. The low concentrations of Aβ*56, which comprise less than 1% of total soluble Aβ in 20 month Tg2576 mice^2^, necessitate taking advantage of the greater sensitivity of biotin-avidin coupled to chemiluminescence compared to colorimetric methods. It is important to beware that some batches of A11 do not detect Aβ*56 (**Fig. S7**).

We surmise, based on comparing banding patterns in our current western blots with those in the literature, that the Aβ*56 entity we describe here overlaps with but may not correspond exactly to the ∼56-kDa entities that have been identified using other protocols. Technical refinements permit more accurate characterizations of Aβ assemblies. For example, the constituents of highly neurotoxic entities in 7PA2 culture medium originally believed to be dimers and trimers^42^ were found upon later investigation also to contain N-terminally extended, non-canonical Aβ^43^. While our current protocol reliably and reproducibly detects Aβ*56, we anticipate that continued, methodological advancements will enable future investigations of Aβ*56 to be conducted more efficiently.

Not all lines of mice produce Aβ*56. The levels and species of Aβ generated in a particular line of mice affect the expression of Aβ*56. For example, Tg2576 mice expressing APP-Swe express Aβ*56, but rTg9191 mice expressing APP-Swe with the V717I London mutation which promotes fibril formation do not^2, 30^. The balance between aggregation processes that do (on-pathway) and do not (off-pathway) culminate in fibrils may influence the production of Aβ*56, as the two pathways may compete for monomers in a common pool.

The structure, spatial distribution, and temporal expression of Aβ oligomers are important determinants of their effects on the brain^2, 44^. Two distinctive spatial and temporal features of Aβ*56 are that it forms before dense-core plaques and is present within but not confined to dense-core plaques^1, 2^. This is significant, because in mice the oligomers that are confined to dense-core plaques do not impair neurological function unless they are burst open and dispersed in the brain^2^. Although the precise structure of Aβ*56 is unknown, some general features may be deduced from binding studies using the polyclonal, conformational antibodies, OC and A11^4, 7^. OC antibodies, which recognize in-register, parallel β-sheets, and to a lesser extent antiparallel β-sheets, do not bind Aβ*56^2^. A11 antibodies, which recognize a variety of conformational epitopes, bind Aβ*56 in mice^1, 20^ (and this paper), dogs^18^, and humans^21, 45^, but the specific epitope(s) recognized is unknown. Because A11 antibodies are polyclonal, more than one A11-reactive variant may be present, complicating the ability to decipher the structural properties of Aβ*56. Currently, there are no monoclonal, conformational antibodies that selectively bind Aβ*56.

In summary, in this paper, we confirmed and extended the characterization of a ∼56-kDa, A11-binding, SDS-stable, water-soluble, non-plaque-related, brain-derived oligomer containing canonical Aβ(1-40), also known as Aβ*56. We showed that its presence correlates with age-related memory loss in Tg2576 mice. We provide detailed information about the techniques, tools, and specimens that enable Aβ*56 to be detected reliably and reproducibly.

### Limitations of the Study

These studies are limited to the Tg2576 mouse model. It is possible that other mouse models express different, SDS-stable, Aβ oligomers, and that Aβ*56 in other mouse models may contain other forms of Aβ, including Aβ(1-42).

Although we did not observe any SDS-stable Aβ oligomers larger than 100 kDa, we cannot exclude their existence because our experimental protocol is optimized to detect oligomers smaller than 100 kDa. Optimized experimental parameters are needed to investigate the existence of larger, SDS-stable oligomers.

While the current studies are limited to mice, it is likely that Aβ*56 is present in humans. One group previously reported detecting higher levels of A11-reactive, ∼56-kDa entities in nasal fluid from humans with AD^21, 38^. We previously described an isolated cluster of ∼56-kDa, 82E1-reactive bands on a western blot of immunoprecipitated Aβ(x-40)/Aβ(x-42) proteins from human cerebrospinal fluid (CSF)^46^. When the immunoprecipitated CSF proteins were probed using 6E10 antibodies, however, non-oligomeric, APP fragments were also detected^46^, illustrating the importance of employing antibodies that specifically detect canonical Aβ. It will be informative to use our current protocols to examine Aβ*56 levels in AD, since an earlier study using 6E10 antibodies reported paradoxically lower levels of Aβ*56 in brain tissue from AD patients^45^.

## Supporting information

SUPPLEMENTARY Liu et al iScience Mar 20 2023

## Author contributions

P.L. conceived the study, designed and performed biochemistry experiments, analyzed data, and wrote the manuscript. I.P.L. and S.L.S. helped design and perform biochemistry experiments, and analyzed data. L.J.K. analyzed archived behavioral data. K.H.A. conceived the study, analyzed data, and wrote the manuscript.

## Acknowledgements

Behavioral studies were supported using funds from NIH R01-NS33249. Biochemistry studies were supported using funds from the N. Bud Grossman Center for Memory Research and Care. The authors thank Dr. Swathy Babu and Dr. Shauna Yuan for contributing Fig. S8.

## Supplementary Figure Legends

Figure S1. The ∼56-kDa, SDS-stable, Aβ(1-40)-containing entities are detected in plaque-bearing Tg2576 mice.

A representative immunoprecipitation (IP)/western blotting (WB) analysis showing that D8Q7I-precipitated entities that migrated at ∼56 kDa (arrow) when fractionated using sodium dodecyl sulfate/polyacrylamide gel electrophoresis (SDS-PAGE) are detected by biotinylated 82E1 (82E1-biotin) antibody directed against the N-terminus of Aβ(1-x) in 14-16 and 21 month Tg2576 mice (+) but not in a 21 month non-transgenic littermate (-). No ∼56-kDa, D8Q7I-precipitated entities are detected when Tg2576 brain extracts were not used (Ab only). Ab, capturing antibody D8Q7I. Syn Aβ_40_, synthetic Aβ(1-40) (5 ng), serving as a positive control for WB. #, monomeric Aβ; $, dimeric Syn Aβ_40_; and *, non-specifically detected entities. kDa, kilo-Daltons.

Figure S2. No ∼56-kDa entities are detected in 82E1-precipitated brain extracts from Tg2576 mice.

**A-B**) A representative immunoprecipitation (IP)/western blotting (WB) analysis showing that ∼56-kDa entities are not detected in the 82E1-precipitated brain extracts by D8Q7I in 8-12 month Tg2576 mice (+) or age-matched non-transgenic littermates (-). Syn Aβ_40_, synthetic Aβ(1-40) (5 ng), serving as a positive control for WB. (A) Light exposure, and (B) heavy exposure. **C-D**) A representative IP/WB analysis showing that ∼56-kDa entities are not detected in the 82E1-precipitated brain extracts by biotinylated D8Q7I (D8Q7I-biotin) in 14-15 and 25 month Tg2576 mice (+) or age-matched non-transgenic littermates (-). No ∼56-kDa, 82E1-precipitated entities are detected when Tg2576 brain extracts were not used (Ab only). Ab, capturing antibody 82E1. (C) short exposure (10 sec), and (D) long exposure (100 sec). #, monomeric Aβ; $, dimeric Aβ; @, trimeric Syn Aβ_40_; and *, non-specifically detected entities. kDa, kilo-Daltons.

Figure S3. Aβ(x-42) is not detected in the ∼56-kDa, SDS-stable entities in Tg2576 mice.

**A**) A representative immunoprecipitation (IP)/western blotting (WB) analysis showing that no obvious D3E10-precipitated entities that migrated at ∼56 kDa in sodium dodecyl sulfate/polyacrylamide gel electrophoresis are detected by biotinylated 6E10 (6E10-biotin) in 8-12 month Tg2576 mice (+) or age-matched non-transgenic littermates (-). Monomeric Aβ (#) is detected in all seven Tg2576 mice but neither of the non-transgenic littermates. No ∼56-kDa, D3E10-precipitated entities are detected when no Tg2576 brain extracts were used (Ab only). No ∼56-kDa entities are detected in Tg2576 mice when generic rabbit immunoglobulin G (RbIgG) was used to precipitate brain extracts (+°, combined brain extracts of the seven Tg2576 mice, each contributing 14.3% of the total proteins), and the presence of a minute level of monomeric Aβ is likely because that they formed non-specific interactions with RbIgG/Sepharose G matrix. Ab, capturing rabbit polyclonal antibody D3E10 that is directed against the C-terminus of Aβ(x-42). Syn AβpE_3-42_, synthetic Aβ_42_ with the 3^rd^ amino acid being a pyroglutamate (30 ng), serving as a negative control for WB; and Syn Aβ_42_, synthetic Aβ(1-42) (5 ng), serving as a positive control for WB. $, dimeric Syn AβpE_3-42_ and Syn Aβ_42_; @, trimeric Syn AβpE_3-42_; and * and vertical line, non-specifically detected entities. **B-C**) A representative IP/WB analysis showing no detection of the ∼56-kDa entities in the 82E1-precipitated brain extracts by D3E10 in 8-12 month Tg2576 mice (+) or age-matched non-transgenic littermates (-). These images were obtained by stripping the detecting antibody D8Q7I of the blots in **Figures S2A** and **S2B** and reprobing with D3E10. Syn Aβ_42_, synthetic Aβ(1-42) (10 ng), serving as a positive control for WB. (B) Light exposure, and (C) heavy exposure. #, monomeric Aβ; $, dimeric Syn Aβ_42_; @, trimeric Syn Aβ_42_; and *, non-specifically detected entities. kDa, kilo-Daltons.

Figure S4. The ∼56-kDa, SDS-stable, Aβ(1-40)-containing entities are detected using anti-Aβ(x-40) antibodies of different sources.

A representative immunoprecipitation (IP)/western blotting (WB) analysis showing that the ∼56-kDa Aβ entities (arrow) and monomeric Aβ (#) are detected by biotinylated mouse monoclonal 6E10 (6E10-biotin) antibody following IP of brain extracts of a 13-month Tg2576 mouse (+) using either rabbit monoclonal D8Q7I antibody or mouse monoclonal HJ2 antibody, both directed against the C-terminus of Aβ(x-40). Following IP using mouse monoclonal HJ7.4 antibody that is directed against the C-terminus of Aβ(x-42), monomeric Aβ (#) but not the ∼56-kDa Aβ entities is detected by biotinylated 6E10. Neither the ∼56-kDa Aβ entities nor monomeric Aβ is detected by biotinylated 6E10 following IP of brain extracts of an age-matched non-transgenic littermate (-) using D8Q7I, HJ2 or HJ7.4. Syn Aβ_40/42_, synthetic Aβ(1-40) and Aβ(1-42) (total of 1, 2, and 4 ng loaded in the three lanes, respectively, in the order from the far left to the far right), serving as a positive control for WB. $, dimeric Syn Aβ_40/42_; @, trimeric Syn Aβ_40/42_; and *, non-specifically detected entities. kDa, kilo-Daltons. M, Precision Plus Protein Standard (Bio-Rad).

Figure S5. The ∼56-kDa, Aβ(1-40)-containing entities in Tg2576 mice are not detected in size-exclusion chromatography fractions without anti-Aβ antibodies.

Representative western blotting (WB) analyses showing the detection of entities in even-numbered size-exclusion chromatography (SEC) fractions (Frac. 34-96) of aqueous brain extracts from 10-11 month Tg2576 mice using horseradish peroxidase (HRP)-conjugated Neutravidin (Neutravidin-HRP; Invitrogen). No ∼56-kDa entities, ∼14-kDa entities, or monomeric Aβ are detected. The four WB images (i.e., Frac. 34-48, 50-64, 66-80, and 82-96) shown were obtained using the same exposure time (100 sec). The WB band patterns and intensities of the input material (5% (w/v) of the brain extracts used for SEC) within each of the four WB images are comparable; a WB of the input is shown in the far left panel. The elution profile of biomolecule standards (Std.) of various sizes is shown above the WB images. kDa, kilo-Daltons. The detecting reagent Neutravidin-HRP was stripped from these blots and then probed with biotinylated D8Q7I to obtain images shown in **Figure 3C**.

Figure S6. Immunopurified Aβ*56 is not detected without anti-Aβ or anti-oligomer antibodies.

A representative western blotting (WB) analysis showing the detection of D8Q7I-purified entities from 15-21 month Tg2576 mice (+) and age-matched non-transgenic littermates (-) using horseradish peroxidase (HRP)-conjugated Neutravidin (Neutravidin-HRP). No Aβ*56 is detectable (exposure time: 2,900 sec). kDa, kilo-Daltons.

Figure S7. The ∼56-kDa entities are detected by some but not all batches of A11, anti-oligomer antibodies.

**A-D)** Representative western blotting (WB) analyses showing the detection of the ∼56-kDa entities (arrow) in 15-21 month Tg2576 mice (+) but not age-matched littermates (-) using anti-oligomer antibody A11 purchased from StressMarq Biosciences (A) and Rockland (B) but not from Invitrogen (currently Thermo Fisher Scientific) (C) or MilliporeSigma (D). kDa, kilo-Daltons.

Figure S8. The ∼56-kDa, Aβ-containing entities are detected in Tg2576 mice by a different, independent research laboratory.

A representative immunoprecipitation (IP)/western blotting (WB) analysis showing that D8Q7I-precipitated entities size-fractionated by sodium dodecyl sulfate/polyacrylamide gel electrophoresis at ∼56 kDa (arrow) are detected by biotinylated 82E1 (82E1-biotin) in five of six randomly selected 16-17 month Tg2576 mice (+) but not in an age-matched, non-transgenic littermate (-). No ∼56-kDa, D8Q7I-precipitated entities are detected when no Tg2576 brain extracts were used (Ab only). No ∼56-kDa entities are detected in Tg2576 mice when generic rabbit immunoglobulin G (RbIgG) was used to precipitate brain extracts (+°, combined brain extracts of the six Tg2576 mice, each contributing 16.7% of the total proteins). The appearance of monomeric Aβ in the RbIgG IP may reflect non-specific binding of Aβ to IgGs and/or matrix. Ab, capturing antibody D8Q7I. Aβ, amyloid-β. Syn Aβ_40_, synthetic Aβ(1-40) (1 ng), serving as a positive control for WB. #, monomeric Aβ; $, dimeric Syn Aβ_40_; and *, non-specifically detected entities. kDa, kilo-Daltons.

## STAR Methods

### Animals

Tg2576 mice^19^ containing the APP_695_ isoform with the Swedish (K670N, M671L) mutation (APP-Swe) in C57B6/SJL, 129S6, or 129S6FVB background were from colonies in the lab of K.H.A. at the University of Minnesota, Twin Cities, Minnesota. All experiments involving mice were performed in full accordance with the guidelines of the Association for Assessment and Accreditation of Laboratory Animal Care and approved (approval #1202A09927) by the Institutional Animal Care and Use Committee at the University of Minnesota. Both male and female mice were used in experiments. Mice were anesthetized with isoflurane and sacrificed by decapitation; brains were immediately harvested, snap-frozen on dry ice, and stored at −80°C. Vital characteristics of the animals used in this study (**Table S1**) were shown.

### Antibodies

Basic information (**Table S2**) and detailed usage (**Table S3**) of all antibodies used in this paper were shown.

### Brain protein extraction

Detergent-free, water-soluble brain protein extracts were prepared using a protocol adapted from a previous publication.^22^ Briefly, each frozen hemi-forebrain free of olfactory bulb and cerebellum was weighed and then transferred to 5 volumes (i.e., 5 mL per g of wet brain tissue) of ice-cold extraction buffer (25 mM tris(hydroxymethyl)aminomethane-hydrochloric acid (Tris-HCl), pH 7.4; 140 mM sodium chloride (NaCl); 3 mM potassium chloride (KCl); 0.1 mM phenylmethylsulfonyl fluoride (PMSF); 0.2 mM 1,10-phenanthroline monohydrate (phen); 0.1% (v/v) protease inhibitor cocktail (MilliporeSigma, Burlington, MA); 0.1% (v/v) phosphatase inhibitor cocktail A (SantaCruz Biotechnology, Dallas, TX); and 0.1% (v/v) phosphatase inhibitor cocktail 2 (MilliporeSigma)). Each piece of brain tissue was homogenized at room temperature under the highest power for 25 strokes using a Dounce homogenizer (PolyScience, Niles, IL). The resulting material was immediately placed on ice for 5 min, then transferred into an ice-cold, sterilized, 1.5-mL microcentrifuge tube, and centrifuged at 16,100 x *g*, 4°C for 90 min. The supernatant was collected and incubated twice with 25 µL of Protein G Sepharose 4 Fast Flow resin (GE healthcare, Piscataway, NJ), each time at 4°C for 1 hr, to deplete endogenous mouse immunoglobulin G (msIgG). Total protein concentrations were determined using the bicinchoninic acid (BCA) assay (Thermo Fisher Scientific, Waltham, MA). Brain protein extracts were then stored at −80°C in multiple aliquots to avoid repeated freeze-and-thaw procedures.

### Size-exclusion chromatography (SEC)

SEC was performed using a Superdex 200 10/300 GL column (GE healthcare) under the control of the BioLogic DuoFlow Chromatography system (Bio-Rad, Hercules, CA).

To calibrate the column, each entity in a biomolecule standard set—eight globular proteins at the molecular weights of 669, 443, 200, 150, 66, 29, 13.4, and 6.5 kDa, and Blue Dextran, >2,000 kDa (MilliporeSigma)—was individually injected at the dose of 200 µg in 250 µL of 0.22 µm-filtered phosphate-buffered saline (PBS, pH 7.4). Biomolecule standards were eluted at room temperature with 0.22 µm-filtered PBS at a flow rate of 0.5 mL/min and a fraction size of 250 µL. The elution chromatogram of each standard was determined by both absorbance at wavelength 280 nm and the BCA assay using a DTX880 Multimode detector (Beckman Coulter, Brea, CA).

To analyze biological samples, detergent-free, water-soluble brain protein extracts of Tg2576 mice at 10-11 months of age prior to the appearance of ThioS-reactive, dense-core plaques in the forebrain^31^ were pooled from 5 mice (**Tables S1 and S3**), each contributing 20% protein mass. The prepared sample was injected at the dose of 2 mg in 250 µL. Proteins were eluted, and the elution chromatogram was determined as described above. Immediately after the elution was complete, the microplate that collected the fractions was stored on ice. Each of the 96 fractions was transferred into a 1.5-mL sterilized microcentrifuge tube that contained 2.6 mM PMSF, 5.2 mM phen, 2.6% (v/v) protease inhibitor cocktail, 2.6% (v/v) phosphatase inhibitor cocktail A, and 2.6% (v/v) phosphatase inhibitor cocktail 2 in 10 µL of PBS. After mixing with an SEC fraction, the final concentrations of the five types of inhibitors were 0.1 mM, 0.2 mM, 0.1% (v/v), 0.1% (v/v), and 0.1% (v/v), respectively. Each inhibitor-containing fraction was evenly split into two parts, snap-frozen on dry ice, and stored at −80°C.

### Immunoprecipitation (IP)

IP was performed according to previously published procedures.^2^ Briefly, brain extracts containing 200-400 µg of total proteins (**Tables S1 and S3**) were incubated with capturing reagents (**Tables S1 and S3**) and Protein G-coated matrix (Protein G Sepharose 4 Fast Flow resin (GE healthcare) or Dynabeads Protein G (Thermo Fisher Scientific); **Tables S1 and S3**) at 4°C for 14-16 hr. Subsequently, the matrix was washed under a gently rotating mode (15 rpm) first in wash buffer 1 (50 mM Tris-HCl, pH 7.4; 300 mM NaCl; 1 mM ethylenediaminetetraacetic acid (EDTA); 0.1% (v/v) polyethylene glycol p-(1,1,3,3-tetramethylbutyl)-phenyl ether (Triton X-100); 0.1 mM PMSF; 0.2 mM phen; 0.1% (v/v) protease inhibitor cocktail; 0.1% (v/v) phosphatase inhibitor cocktail A; and 0.1% (v/v) phosphatase inhibitor cocktail 2) at 4°C for 5 min, and then in wash buffer 2 (50 mM Tris-HCl, pH 7.4; 150 mM NaCl; 1 mM EDTA; 0.1% (v/v) Triton X-100; 0.1 mM PMSF; 0.2 mM phen; 0.1% (v/v) protease inhibitor cocktail; 0.1% (v/v) phosphatase inhibitor cocktail A; and 0.1% (v/v) phosphatase inhibitor cocktail 2) at 4°C for 5 min.

### Western blotting (WB)

WB was performed based on previously published procedures.^2, 28^ Detailed usage of brain protein extracts and detecting reagents are listed in Table S3.

Briefly, eluted materials from IP (**Figs. 1B, 2B, 4, S1-S4, and S8**), SEC-fractionated brain extracts (**Figs. 3A, 3C, and S5**), immunopurified entities (**Figs. 5A–5C and S6**), and brain extracts (**Figs. 3A, 3C, and S7**) were electrophoretically separated by SDS-PAGE. To prepare samples for SDS-PAGE, immunoprecipitated materials (**Figs. 1B, 2B, 4, S1-S4, and S8**) were eluted by agitatedly heating the matrix in 4x Laemmli sample buffer (Bio-Rad; plus 1.42 M β-mercaptoethanol) at 1,250 rpm, 95°C for 10 min. SEC-fractionated brain extracts (**Figs. 3A, 3C, and S5**), immunopurified entities (**Figs. 5A and 5B**), or brain extracts (**Figs. 3A and 3C**) were mixed with 4x Laemmli sample buffer plus 1.42 M β-mercaptoethanol at 1,250 rpm, 95°C for 5 min. Immunopurified entities (**Fig. 5C and S6**) or brain extracts (**Fig. S7**) were mixed with 4x Laemmli sample buffer without β-mercaptoethanol or heating. Electrophoresis was performed at room temperature in running buffer 1 (Cathode buffer: 100 mM Tris-HCl, pH 8.25; 100 mM tricine; 0.1% (w/v) SDS. Anode buffer: 200 mM Tris-HCl, pH 8.9) under a constant voltage of 80 V for 150 min when using 10-20% Criterion Tris-tricine gels (Bio-Rad; **Figs. 2B, 4, S2A, S2B, S3, and S4**) or in running buffer 2 (100 mM Tris-HCl, pH 8.3; 100 mM tricine, 0.1% (w/v) SDS) under a constant voltage of 125 V for 90 min when using Novex 10-20% tricine gels (Thermo Fisher Scientific; **Figs. 1B, 3A, 3C, 5, S1, S2C, S2D, S5-S7, and S8**).

The size-fractionated proteins were then transferred to 0.2-µm (pore size) nitrocellulose membranes (Bio-Rad) in transfer buffer 1 (25 mM Tris-HCl, 192 mM glycine, 10% (v/v) methanol) at 4°C under a constant current of 0.4 A for 4 hr when using 10-20% Criterion Tris-tricine gels or in transfer buffer 2 (25 mM Bicine, 25 mM Bis-Tris, 1 mM EDTA, pH 7.2; 0.1% (v/v) NuPAGE antioxidant (Thermo Fisher Scientific); 10% (v/v) methanol) at room temperature or 4°C under a constant voltage of 25 V for 2 hr when using Novex 10-20% tricine gels.

Next, each protein-blotted membrane was transferred to 50 mL of PBS of room temperature, and the membrane was subjected to microwave heating under full power first for 25 sec and then for 15 sec with a 4-min cooling period after each episode of heating.

Following heat-induced antigen retrieval, membranes were incubated with blocking buffers at 70 rpm, room temperature for 1 hr to block non-specific binding of the detecting reagents. For WB analyses using A11 antibodies (**Figs. 5B, 5C, S6, and S7**), blocking buffer 1 (10 mM Tris-HCl, pH 7.4; 200 mM NaCl; 0.01% (v/v) polyoxyethylene (20) sorbitan monolaurate (Tween-20); 5% (w/v) Carnation instant non-fat dry milk powder) of room temperature was used; and for the rest WB analyses, blocking buffer 2 (10 mM Tris-HCl, pH 7.4; 200 mM NaCl; 0.1% (v/v) Tween-20; 5% (w/v) bovine serum albumin (MilliPoreSigma)) was used.

Next, primary antibodies were then added directly to blocking buffer (see **Tables S2 and S3** for the use of detecting antibodies in detail), and membranes were then incubated at 70 rpm, 4°C for 14-16 hr (For **Figs. S5 and S6**, membranes were incubated directly with blocking buffer at 70 rpm, 4°C for 14-16 hr.).

Following primary antibody incubation, membranes were washed with wash buffers five times, each time at 80 rpm, room temperature for 5 min. For WB analyses using A11 antibodies (**Figs. 5B, 5C, and S7**) as well as of Figure S6, wash buffer 3 (10 mM Tris-HCl, pH 7.4; 200 mM NaCl; 0.01% (v/v) Tween-20) of room temperature was used; and for the rest WB analyses, wash buffer 4 (10 mM Tris-HCl, pH 7.4; 200 mM NaCl; 0.1% (v/v) Tween-20) of room temperature was used.

For WB analyses probed with non-biotinylated primary antibodies (**Figs. S2A, S2B, S3B, S3C, and S7**), membranes were then incubated with biotinylated Biotin-SP (long spacer) AffiniPure donkey-anti-rabbit IgG (H+L) (Jackson ImmunoResearch Laboratories, West Grove, PA; diluted 1:50,000 in either blocking buffer 1 of room temperature for A11 (**Fig. S7**) or blocking buffer 2 of room temperature for D8Q7I (**Figs. S2A and S2B**) or D3E10 (**Figs. S3B and S3C**)) at 70 rpm, room temperature for 1 hr. Following the incubation with the secondary antibody, membranes were washed in wash buffers as described above. Next, membranes were first incubated with horseradish peroxidase (HRP)-conjugated Neutravidin (Neutravidin-HRP; Thermo Fisher Scientific; diluted 1:5,000 in either wash buffer 3 of room temperature (**Fig. S7**) or wash buffer 4 of room temperature (**Figs. S2A, S2B, S3B, and S3C**)) at 70 rpm, room temperature for 10 (**Figs. S2A, S2B, S3B, and S3C**) or 30 (**Fig. S7**) min, and then washed in wash buffers as described above.

For WB analyses probed with biotinylated primary antibodies (**Figs. 1B, 2B, 3A, 3C, 4, 5, S1, S2C, S2D, S3A, S4, and S8**) or no primary antibody (**Figs. S5 and S6**), membranes were then directly incubated with Neutravidin-HRP (diluted 1:5,000 in either wash buffer 3 of room temperature (**Figs. 5B and 5C** for biotinylated A11, and **Fig. S6** for Neutravidin-HRP) or wash buffer 4 of room temperature (**Figs. 1B, 2B, 3A, 3C, 4, 5A, S1, S2C, S2D, S3A, S4, S5, and S8**)) at 70 rpm, room temperature for 10 (**Figs. 1B, 2B, 4, S3A, and S4**) or 30 (**Figs. 3A, 3C, 5, S1, S2C, S2D, S5, S6, and S8**) min. Next, membranes were washed wash buffers as described above.

Membranes were then incubated with the SuperSignal West Femto Maximum Sensitivity Substrate (Thermo Fisher Scientific) at 200 rpm, room temperature for 5 min. Chemiluminescence signals were developed using the Kodak Scientific Imaging film X-OMAT Blue XB (PerkinElmer Life Sciences, Waltham, MA). Intensities of immunoreactive protein bands were determined densitometrically using Optiquant version 03.00 (Packard Instrument, Fallbrook, CA). Alternatively, signals were revealed using the ChemiDoc MP Imaging System (Bio-Rad). Image Lab version 6.1 (Bio-Rad) was used to determine protein band intensities.

To determine whether the ∼56-kDa, Aβ-containing entity was present in each studied animal, the band total intensity (i.e., the product of mean intensity per pixel and numbers of pixels in the designated band area) of the ∼56-kDa, Aβ-containing entity was first measured. Next, the mean and standard deviation (SD) of background intensities were determined from the measured total intensities of four areas, each of which had an equal size to the size of the designated band area of the ∼56-kDa, Aβ-containing entity. The four areas were sampled at 1) the ∼56-kDa in the lane containing immunoprecipitated entities from non-transgenic mice, 2) the ∼4.5-kDa (corresponding to the migrating area of monomeric Aβ) in the lane containing immunoprecipitated entities from non-transgenic mice, 3) the ∼56-kDa in the lane containing only capturing antibodies but no immunoprecipitated entities from mice, and 4) the ∼4.5-kDa in the lane containing only capturing antibodies but no immunoprecipitated entities from mice. The ∼56-kDa, Aβ-containing entity was considered detected (i.e., present) in the brain of an animal if its band total intensity is equal to or higher than three SDs above the mean background intensity.

The experimental settings for each (IP)/WB reaction are shown in Table S3.

### Denaturant treatment

Three hundred and seventy-five µg of mouse brain extracts (**Tables S1 and S3**) were completely dried but not over-dried at room temperature using a speed vacuum concentrator (Eppendorf, Hamburg, Germany). The resulting materials were first gently and fully resuspended in 70 µL of 8 M urea (in 20 mM Tris-HCl, pH 8.0), 6 M GuHCl (in 20 mM Tris-HCl, pH 8.0), 100% (v/v) HFIP, or TBS (50 mM Tris-HCl, pH 7.4; 150 mM NaCl; serving as a positive control for the detection of the ∼56-kDa, Aβ-containing entity), and then agitated at 4°C, 1,200 rpm for 16-18 hr. Brain extracts of age-matched, non-transgenic mice similarly treated with TBS served as a negative control for the detection of the ∼56-kDa, Aβ-containing entity. Following the treatments, the samples were dried using the speed vacuum concentrator at room temperature for 15 min to evaporate the solvents, 1.4 mL of TBS (containing 0.1 mM PMSF, 0.2 mM phen, 0.1% (v/v) protease inhibitor cocktail, 0.1% (v/v) phosphatase inhibitor cocktail A, and 0.1% (v/v) phosphatase inhibitor cocktail 2) was added to each sample, and the samples were gently resuspended until they were fully homogenized. The resulting materials were subjected to IP as described in the section “*Immunoprecipitation (IP)*.”

### Antibody immobilization on Dynabeads Protein G

The antibody immobilization was carried out as previously described.^46, 47^ Briefly, 62 µg of the D8Q7I antibody was incubated in 1 mL of TBS with Dynabeads Protein G matrix from 1 mL of fully resuspended slurry (settled bead volume: ∼100 µL) under a gently rotating mode (15 rpm) at 4°C for 14-16 hr. The antibody was covalently linked to the matrix by incubating with 1 mL of 70 mM dimethyl pimelimidate (Thermo Fisher Scientific; dissolved in triethanolamine buffer solution (MilliporeSigma) that contains 200 mM triethanolamine and pH adjusted to 9.0) solution under a gently rotating mode (15 rpm) at room temperature for 15 min. The crosslinking reaction was then quenched by incubating the antibody-matrix complex with 1 mL of 10% (v/v) ethanolamine (MilliporeSigma; diluted 100% (v/v) stock solution 10-fold in distilled deionized water) solution under a gently rotating mode (15 rpm) at room temperature for 15 min. Prior to use, beads were pre-treated twice with elution buffer 1 (100 mM glycine-HCl, pH 2.9; 1 M urea) under a gently rotating mode (15 rpm) at room temperature for 5 min to wash off any antibody not covalently linked to beads. The antibody-matrix complex was stored at 4°C in TBS containing 0.05% (w/v) sodium azide and inhibitors (0.1 mM PMSF, 0.2 mM phen, 0.1% (v/v) protease inhibitor cocktail, 0.1% (v/v) phosphatase inhibitor cocktail A, and 0.1% (v/v) phosphatase inhibitor cocktail 2).

### Immunoaffinity purification

Brain protein extracts (**Tables S1 and S3**) of 15-21 month Tg2576 or age-matched non-transgenic mice were incubated with D8Q7I-bound Dynabeads Protein G matrix (**Table S3**) under a gently rotating mode (15 rpm) at 4°C for 20 hr. The immunocomplex-matrix was then washed twice with 1 mL of wash buffer 5 (TBS plus 1% (w/v) *n*-Octyl *β*-D-thioglucopyranoside (OTG)) under a gently rotating mode (15 rpm) at 4°C for 5 min. Immunoprecipitated entities were eluted three times using 50 μL of elution buffer 2 (Pierce IgG Elution Buffer, pH ∼3 (Thermo Fisher Scientific); plus 1% (w/v) OTG) by agitating at 1,200 rpm, 25°C for 5 min. Each time immediately after the elution, the pH of the eluted material was neutralized to pH between 7 and 8 using a neutralization solution (1 M Tris-base (pH ∼10.5)). The eluted materials were stored at −80°C.

### Behavioral testing

We analyzed archived data generated as described in Westerman *et al*.^32^ Briefly, we tested spatial reference learning and memory using a version of the conventional Morris water maze.^48^ The water maze was a circular 1- or 1.2-m pool filled with water at 25-27°C and made opaque by the addition of nontoxic white paint. The pool was placed amid fixed spatial cues consisting of boldly patterned curtains and shelves containing distinct objects. Mice first underwent visible platform training for 3 consecutive days (eight trials per day), swimming to a raised platform (a square surface 12 x 12 cm^2^) marked with a black and white striped pole. Hidden platform training was conducted over 9 consecutive days (four trials per day), wherein mice were allowed to search for a platform submerged 1.5 cm beneath the surface of the water. At the beginning of the 4^th^, 7^th^, and 10^th^ day of hidden platform training, a probe trial was conducted in which the platform was removed from the pool, and mice were allowed to search for the platform for 60 sec. All trials were monitored by a camera mounted directly above the pool, and were recorded and analyzed using a computerized tracking system (HVS image, Hampton UK). Further analysis was done using Wintrack (kindly provided by Dr. David Wolfer, University of Zurich, Switzerland). The mean probe score (MPS) is the arithmetic average of the %-time spent in the target quadrant in the three probe trials.

### Key resource tables may be found in the Supplemental Information

Table S1 Mice used to detect the ∼56-kDa, Aβ-containing entities

Table S2 Antibodies used in this study

Table S3 Mice and reagents used in immunoprecipitation, immunoaffinity purification, and western blotting

