## SUPPLEMENTARY Liu et al iScience Mar 20 2023 for "Aβ*56 is a stable oligomer that correlates with age-related memory loss in Tg2576 mice"

### FIGURES and TABLES

Liu *et al*

**Figure S1. The ~56-kDa, SDS-stable, Aβ(1-40)-containing entities are detected in plaque-bearing Tg2576 mice.**

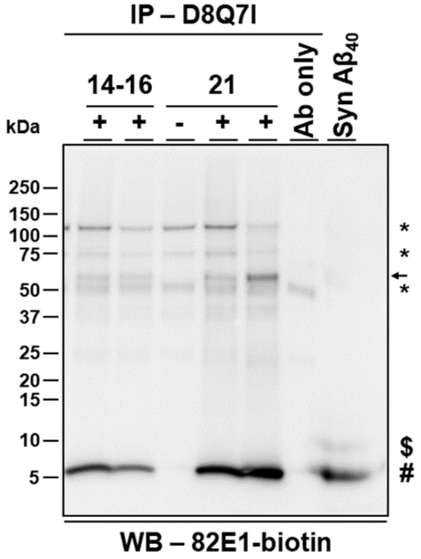

**Figure S2. No ~56-kDa entities are detected in 82E1-precipitated brain extracts from Tg2576 mice**.

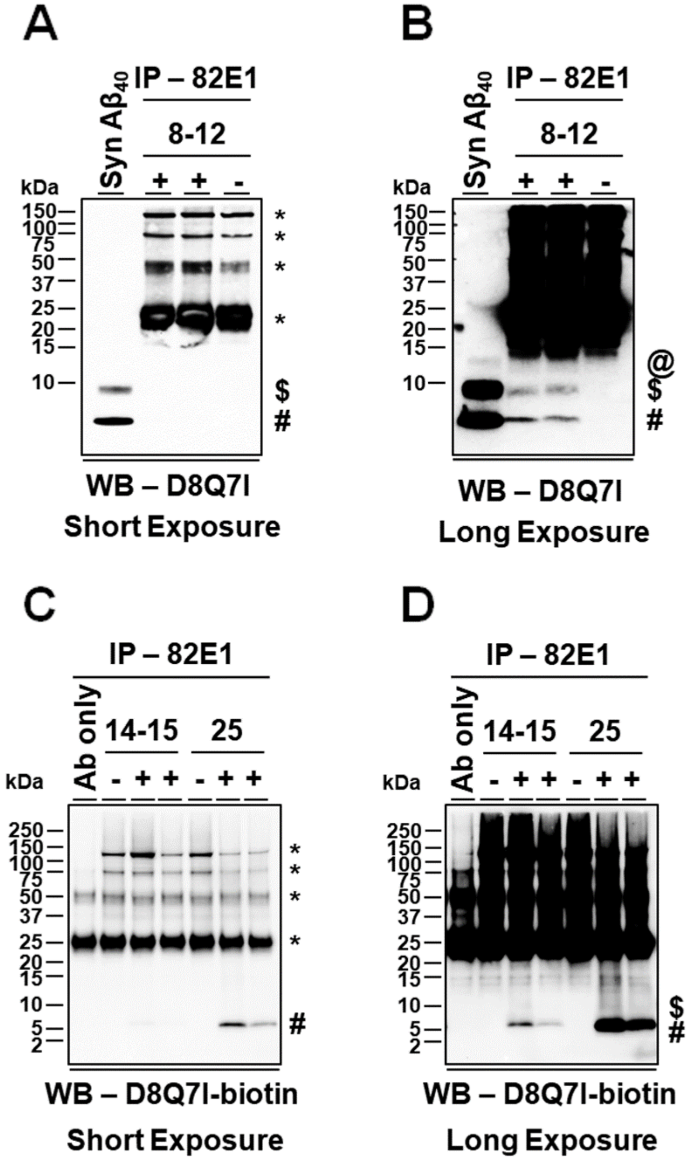

**Figure S3. Aβ(x-42) is not detected in the ~56-kDa, SDS-stable entities in Tg2576 mice.**

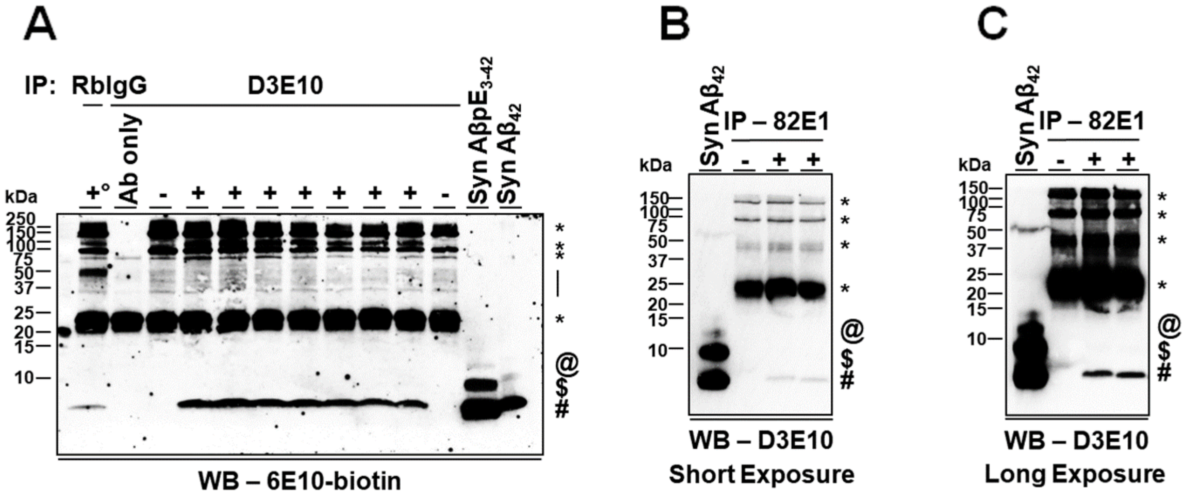

**Figure S4. The ~56-kDa, SDS-stable, Aβ(1-40)-containing entities are detected using anti-Aβ(x-40) antibodies from different sources**.

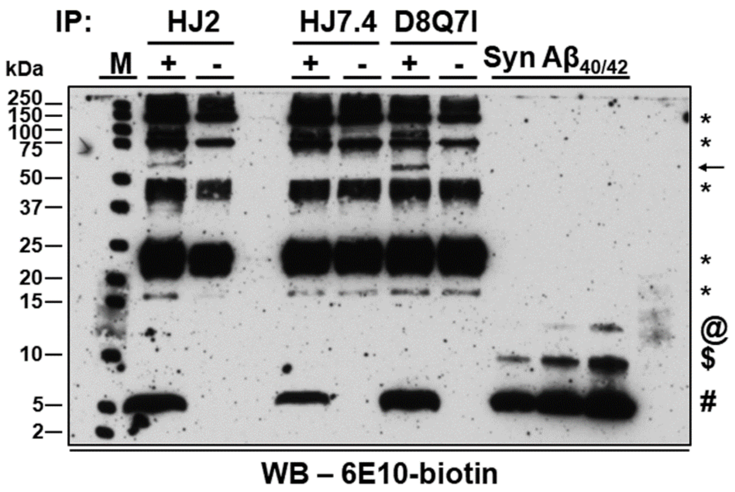

**Figure S5. The ~56-kDa, Aβ(1-40)-containing entities in Tg2576 mice are not detected in SEC fractions without anti-Aβ antibodies.**

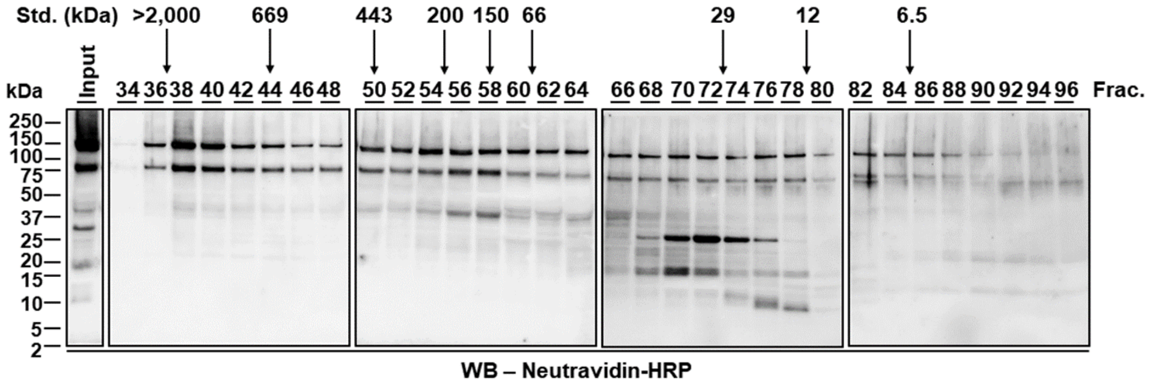

**Figure S6. Immunopurified Aβ*56 is not detected without anti-Aβ or anti-oligomer antibodies.**

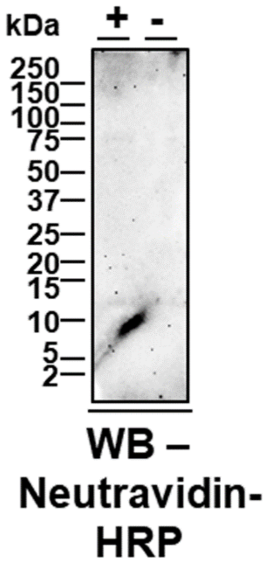

**Figure S7. The ~56-kDa entities are detected by some but not all batches of A11, anti-oligomer antibodies.**

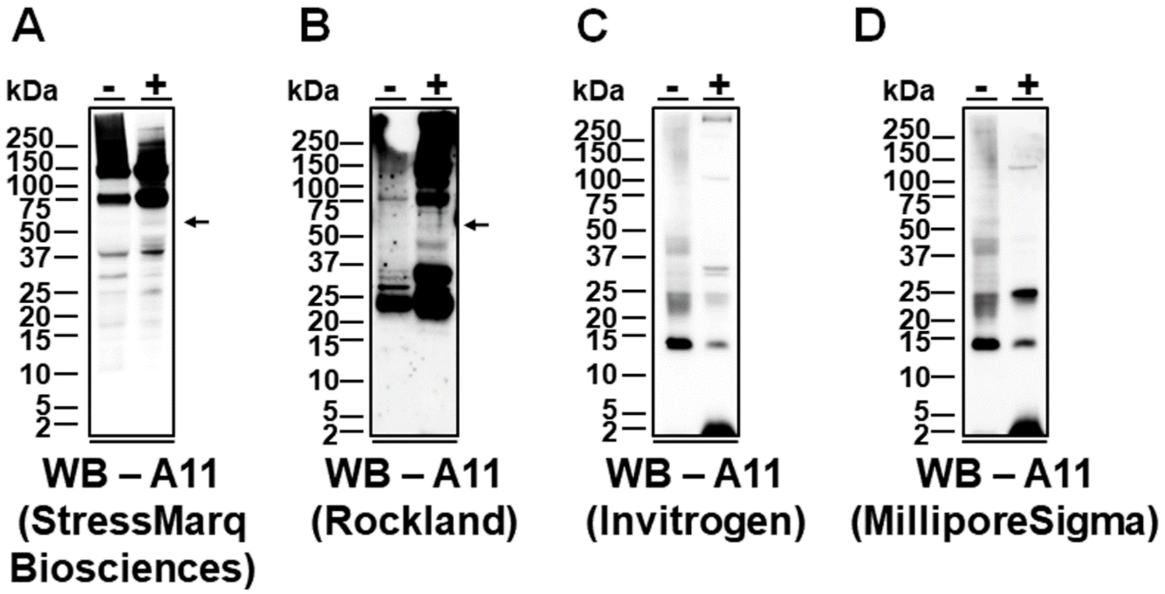

**Figure S8. The ~56-kDa, Aβ-containing entities are detected in Tg2576 mice by a different, independent research laboratory.**

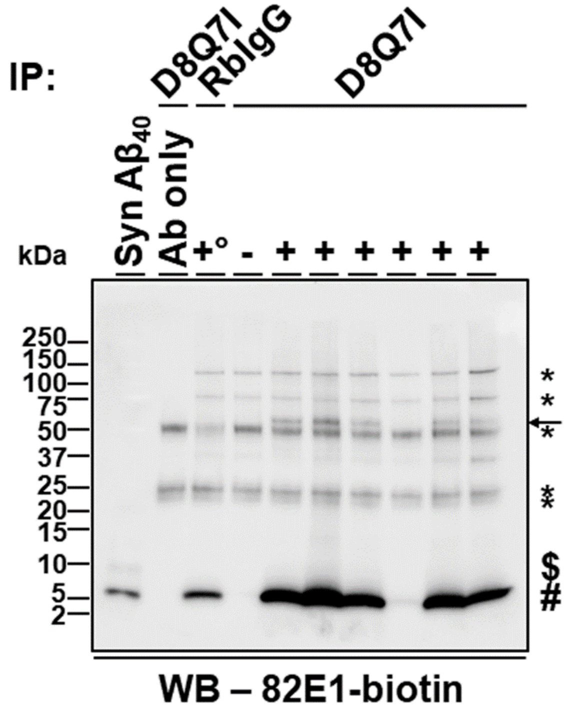

**Table S1 Mice used to detect the ~56-kDa, Aβ-containing entities**

| Line | ID | Tg[a] | Background | Age (Mo.)[b] | DOB[c] | Gender[d] | ~56-kDa[e] |
|---|---|---|---|---|---|---|---|
| Tg2576 | 1 | - | 129S6 | 2.0 | 03/16/2002 | M | N |
| Tg2576 | 2 | + | 129S6 | 2.0 | 03/16/2002 | M | N |
| Tg2576 | 3 | + | 129S6 | 2.0 | 03/16/2002 | M | N |
| Tg2576 | 4 | + | 129S6 | 2.0 | 03/16/2002 | M | N |
| Tg2576 | 5 | - | 129S6 | 3.0 | 08/18/2012 | F | N |
| Tg2576 | 6 | + | 129S6 | 3.1 | 08/17/2012 | F | N |
| Tg2576 | 7 | + | 129S6 | 3.1 | 08/17/2012 | F | N |
| Tg2576 | 8 | + | 129S6 | 3.1 | 08/17/2012 | F | N |
| Tg2576 | 9 | - | 129S6 | 3.1 | 08/18/2012 | F | N |
| Tg2576 | 10 | + | B6S | 4.0 | 09/24/2007 | F | N |
| Tg2576 | 11 | + | B6S | 4.0 | 09/24/2007 | M | N |
| Tg2576 | 12 | + | B6S | 4.0 | 09/24/2007 | M | N |
| Tg2576 | 13 | + | B6S | 5.3 | 02/04/2004 | M | N |
| Tg2576 | 14 | + | B6S | 5.3 | 02/06/2004 | F | N |
| Tg2576 | 15 | + | B6S | 5.3 | 02/06/2004 | F | N |
| Tg2576 | 16 | + | 129S6/FVB | 6.6 | 08/09/2005 | F | N |
| Tg2576 | 17 | + | 129S6/FVB | 6.6 | 08/09/2005 | F | N |
| Tg2576 | 18 | + | 129S6/FVB | 6.6 | 08/09/2005 | F | N |
| Tg2576 | 19 | - | 129S6/FVB | 6.8 | 09/18/2015 | F | N |
| Tg2576 | 20 | + | 129S6/FVB | 6.8 | 08/09/2005 | M | N |
| Tg2576 | 21 | + | 129S6/FVB | 6.8 | 08/09/2005 | M | N |
| Tg2576 | 22 | + | 129S6/FVB | 6.8 | 08/09/2005 | M | N |
| Tg2576 | 23 | - | B6S | 7.0 | 12/20/2006 | F | N |
| Tg2576 | 24 | + | B6S | 7.0 | 12/20/2006 | F | Y |
| Tg2576 | 25 | + | 129S6/FVB | 7.1 | 09/08/2015 | F | N |
| Tg2576 | 26 | + | 129S6/FVB | 7.1 | 09/08/2015 | F | N |
| Tg2576 | 27 | - | 129S6/FVB | 7.1 | 09/08/2015 | F | N |
| Tg2576 | 28 | + | B6S | 7.1 | 12/31/2006 | M | N |
| Tg2576 | 29 | - | B6S | 7.1 | 01/05/2004 | F | N |
| Tg2576 | 30 | + | 129S6/FVB | 7.2 | 09/06/2015 | F | N |
| Tg2576 | 31 | + | 129S6/FVB | 7.2 | 09/07/2015 | F | N |
| Tg2576 | 32 | + | 129S6/FVB | 7.2 | 09/07/2015 | F | Y |
| Tg2576 | 33 | + | B6S | 7.2 | 12/27/2006 | F | N |
| Tg2576 | 34 | + | B6S | 7.2 | 01/05/2004 | F | N |
| Tg2576 | 35 | + | B6S | 7.2 | 01/05/2004 | M | N |
| Tg2576 | 36 | + | B6S | 7.2 | 01/05/2004 | F | N |
| Tg2576 | 37 | + | B6S | 7.2 | 01/05/2004 | M | N |
| Tg2576 | 38 | + | B6S | 7.2 | 01/05/2004 | M | N |
| Tg2576 | 39 | - | B6S | 7.2 | 01/10/2004 | F | N |
| Tg2576 | 40 | + | B6S | 7.4 | 12/20/2006 | F | N |
| Tg2576 | 41 | + | B6S | 7.4 | 12/20/2006 | F | N |
| Tg2576 | 42 | + | B6S | 7.4 | 12/20/2006 | F | N |
| Tg2576 | 43 | + | B6S | 7.4 | 12/20/2006 | F | N |

| Line | ID | Tg[a] | Background | Age (Mo.)[b] | DOB[c] | Gender[d] | ~56-kDa[e] |
|---|---|---|---|---|---|---|---|
| Tg2576 | 44 | - | B6S | 7.4 | 02/09/2004 | M | N |
| Tg2576 | 45 | + | 129S6 | 8.5 | 07/10/2003 | F | N |
| Tg2576 | 46 | + | 129S6 | 8.5 | 07/10/2003 | F | Y |
| Tg2576 | 47 | + | B6S | 8.6 | 09/13/2005 | M | Y |
| Tg2576 | 48 | + | B6S | 8.6 | 09/13/2005 | M | Y |
| Tg2576 | 49 | + | 129S6 | 8.6 | 07/07/2003 | F | Y |
| Tg2576 | 50 | + | 129S6 | 8.6 | 07/17/2003 | M | N |
| Tg2576 | 51 | + | 129S6 | 8.6 | 07/17/2003 | M | N |
| Tg2576 | 52 | + | 129S6 | 8.6 | 07/17/2003 | M | N |
| Tg2576 | 53 | + | 129S6 | 8.6 | 07/17/2003 | M | N |
| Tg2576 | 54 | + | B6S | 8.7 | 11/09/2007 | F | Y |
| Tg2576 | 55 | + | B6S | 8.7 | 11/09/2007 | F | N |
| Tg2576 | 56 | - | B6S | 8.7 | 11/09/2007 | M | N |
| Tg2576 | 57 | + | B6S | 8.7 | 11/09/2007 | M | N |
| Tg2576 | 58 | + | B6S | 8.7 | 03/08/2012 | F | N |
| Tg2576 | 59 | + | B6S | 8.8 | 04/14/2009 | M | N |
| Tg2576 | 60 | + | B6S | 8.8 | 03/05/2012 | F | N |
| Tg2576 | 61 | + | B6S | 8.8 | 03/08/2012 | M | N |
| Tg2576 | 62 | + | 129S6 | 8.8 | 11/22/1999 | F | N |
| Tg2576 | 63 | + | 129S6 | 8.9 | 06/27/2003 | F | N |
| Tg2576 | 64 | + | 129S6 | 8.9 | 06/27/2003 | M | N |
| Tg2576 | 65 | + | 129S6 | 8.9 | 07/08/2003 | F | N |
| Tg2576 | 66 | + | 129S6 | 8.9 | 07/08/2003 | F | N |
| Tg2576 | 67 | + | 129S6 | 8.9 | 07/08/2003 | F | N |
| Tg2576 | 68 | + | 129S6 | 8.9 | 07/08/2003 | M | N |
| Tg2576 | 69 | + | 129S6 | 8.9 | 07/08/2003 | M | N |
| Tg2576 | 70 | + | 129S6 | 8.9 | 07/10/2003 | M | N |
| Tg2576 | 71 | + | 129S6 | 9.2 | 06/19/2003 | F | N |
| Tg2576 | 72 | + | 129S6 | 9.2 | 06/19/2003 | M | N |
| Tg2576 | 73 | + | B6S | 9.7 | 03/07/2012 | F | N |
| Tg2576 | 74 | + | B6S | 9.7 | 03/07/2012 | F | N |
| Tg2576 | 75 | + | B6S | 9.7 | 03/05/2012 | M | N |
| Tg2576 | 76 | + | 129S6 | 9.7 | 05/28/2003 | F | Y |
| Tg2576 | 77 | + | 129S6 | 9.7 | 06/03/2003 | F | N |
| Tg2576 | 78 | + | B6S | 9.8 | 03/08/2012 | M | Y |
| Tg2576 | 79 | + | B6S | 9.8 | 03/08/2012 | F | N |
| Tg2576 | 80 | + | 129S6 | 9.8 | 05/30/2003 | F | Y |
| Tg2576 | 81 | + | 129S6 | 9.8 | 05/30/2003 | M | Y |
| Tg2576 | 82 | + | 129S6 | 9.8 | 06/01/2003 | F | N |
| Tg2576 | 83 | + | 129S6 | 9.8 | 06/01/2003 | F | N |
| Tg2576 | 84 | + | 129S6 | 9.8 | 06/01/2003 | M | N |
| Tg2576 | 85 | + | B6S | 9.9 | 10/03/2007 | F | Y |
| Tg2576 | 86 | - | B6S | 9.9 | 11/09/2007 | F | N |
| Tg2576 | 87 | + | B6S | 9.9 | 03/07/2012 | M | Y |

| Line | ID | Tg[a] | Background | Age (Mo.)[b] | DOB[c] | Gender[d] | ~56-kDa[e] |
|---|---|---|---|---|---|---|---|
| Tg2576 | 88 | + | 129S6 | 9.9 | 05/28/2003 | M | N |
| Tg2576 | 89 | - | B6S | 10.0 | 10/02/2007 | F | N |
| Tg2576 | 90 | + | B6S | 10.0 | 10/02/2007 | F | N |
| Tg2576 | 91 | + | B6S | 10.0 | 10/02/2007 | F | Y |
| Tg2576 | 92 | + | B6S | 10.0 | 10/02/2007 | F | N |
| Tg2576 | 93 | - | B6S | 10.0 | 10/02/2007 | F | N |
| Tg2576 | 94 | - | B6S | 10.2 | 09/13/2005 | M | N |
| Tg2576 | 95 | + | 129S6 | 10.2 | 12/26/2007 | M | N |
| Tg2576 | 96 | - | 129S6 | 10.2 | 12/26/2007 | F | N |
| Tg2576 | 97 | - | 129S6 | 10.2 | 12/26/2007 | F | N |
| Tg2576 | 98 | + | 129S6 | 10.2 | 12/26/2007 | M | N |
| Tg2576 | 99 | + | 129S6 | 10.2 | 12/26/2007 | M | Y |
| Tg2576 | 100 | + | B6S | 10.2 | 09/13/2005 | F | N |
| Tg2576 | 101 | + | B6S | 10.2 | 09/13/2005 | F | N |
| Tg2576 | 102 | + | B6S | 10.2 | 09/13/2005 | F | N |
| Tg2576 | 103 | + | 129S6 | 10.2 | 02/27/2001 | M | N |
| Tg2576 | 104 | + | 129S6 | 10.2 | 03/04/2001 | M | N |
| Tg2576 | 105 | + | 129S6 | 10.2 | 03/04/2001 | M | N |
| Tg2576 | 106 | + | 129S6 | 10.2 | 03/04/2001 | M | Y |
| Tg2576 | 107 | + | B6S | 10.3 | 09/13/2005 | M | N |
| Tg2576 | 108 | + | B6S | 10.3 | 09/13/2005 | M | N |
| Tg2576 | 109 | + | B6S | 10.3 | 09/15/2005 | M | N |
| Tg2576 | 110 | + | B6S | 10.5 | 08/07/2006 | M | N |
| Tg2576 | 111 | + | B6S | 10.6 | 07/15/2001 | M | N |
| Tg2576 | 112 | + | B6S | 10.6 | 07/15/2001 | M | N |
| Tg2576 | 113 | + | B6S | 10.6 | 07/15/2001 | F | Y |
| Tg2576 | 114 | + | B6S | 10.6 | 07/15/2001 | F | N |
| Tg2576 | 115 | + | B6S | 10.6 | 07/15/2001 | F | N |
| Tg2576 | 116 | + | B6S | 10.6 | 07/15/2001 | M | Y |
| Tg2576 | 117 | - | B6S | 10.7 | 09/17/2007 | F | N |
| Tg2576 | 118 | + | B6S | 10.7 | 09/17/2007 | F | N |
| Tg2576 | 119 | + | 129S6/FVB | 10.8 | 09/18/2015 | F | N |
| Tg2576 | 120 | + | 129S6/FVB | 10.8 | 09/18/2015 | F | Y |
| Tg2576 | 121 | - | 129S6/FVB | 10.8 | 06/23/2005 | M | N |
| Tg2576 | 122 | - | 129S6/FVB | 10.8 | 06/23/2005 | F | N |
| Tg2576 | 123 | - | 129S6/FVB | 10.8 | 06/23/2005 | F | N |
| Tg2576 | 124 | + | B6S | 10.9 | 06/30/2008 | F | N |
| Tg2576 | 125 | + | 129S6/FVB | 10.9 | 02/14/2016 | F | Y |
| Tg2576 | 126 | + | 129S6/FVB | 10.9 | 02/14/2016 | F | Y |
| Tg2576 | 127 | + | 129S6/FVB | 10.9 | 02/14/2016 | F | Y |
| Tg2576 | 128 | - | 129S6/FVB | 10.9 | 02/14/2016 | F | N |
| Tg2576 | 129 | + | 129S6/FVB | 10.9 | 06/23/2005 | F | Y |
| Tg2576 | 130 | + | 129S6/FVB | 10.9 | 06/23/2005 | M | Y |
| Tg2576 | 131 | + | 129S6/FVB | 10.9 | 06/23/2005 | M | N |

| Line | ID | Tg[a] | Background | Age (Mo.)[b] | DOB[c] | Gender[d] | ~56-kDa[e] |
|---|---|---|---|---|---|---|---|
| Tg2576 | 132 | + | 129S6/FVB | 11.0 | 02/10/2016 | F | N |
| Tg2576 | 133 | + | 129S6/FVB | 11.0 | 02/10/2016 | F | Y |
| Tg2576 | 134 | + | 129S6/FVB | 11.0 | 02/10/2016 | F | Y |
| Tg2576 | 135 | - | 129S6/FVB | 11.0 | 02/20/2005 | F | N |
| Tg2576 | 136 | + | 129S6/FVB | 11.0 | 02/20/2005 | F | N |
| Tg2576 | 137 | + | 129S6/FVB | 11.0 | 02/20/2005 | F | N |
| Tg2576 | 138 | + | 129S6/FVB | 11.0 | 02/25/2005 | F | Y |
| Tg2576 | 139 | + | 129S6/FVB | 11.0 | 02/25/2005 | F | Y |
| Tg2576 | 140 | + | 129S6/FVB | 11.0 | 02/25/2005 | M | Y |
| Tg2576 | 141 | + | 129S6/FVB | 11.0 | 02/23/2005 | F | Y |
| Tg2576 | 142 | + | 129S6/FVB | 11.0 | 02/23/2005 | M | Y |
| Tg2576 | 143 | + | 129S6/FVB | 11.0 | 02/23/2005 | M | Y |
| Tg2576 | 144 | + | 129S6/FVB | 11.0 | 06/21/2005 | M | N |
| Tg2576 | 145 | - | 129S6/FVB | 11.1 | 09/08/2015 | F | N |
| Tg2576 | 146 | + | 129S6/FVB | 11.1 | 09/08/2015 | F | N |
| Tg2576 | 147 | + | 129S6/FVB | 11.1 | 09/08/2015 | F | Y |
| Tg2576 | 148 | - | 129S6/FVB | 11.1 | 06/23/2005 | F | N |
| Tg2576 | 149 | + | 129S6/FVB | 11.1 | 01/27/2003 | M | N |
| Tg2576 | 150 | + | 129S6/FVB | 11.1 | 01/27/2003 | M | N |
| Tg2576 | 151 | + | 129S6/FVB | 11.2 | 09/07/2015 | F | N |
| Tg2576 | 152 | + | 129S6/FVB | 11.2 | 09/07/2015 | F | N |
| Tg2576 | 153 | + | 129S6/FVB | 11.2 | 09/07/2015 | F | Y |
| Tg2576 | 154 | + | 129S6/FVB | 11.2 | 09/07/2015 | F | N |
| Tg2576 | 155 | + | 129S6/FVB | 11.2 | 02/03/2016 | F | Y |
| Tg2576 | 156 | + | 129S6/FVB | 11.2 | 02/03/2016 | F | N |
| Tg2576 | 157 | - | 129S6/FVB | 11.2 | 02/03/2016 | F | N |
| Tg2576 | 158 | - | 129S6/FVB | 11.2 | 02/03/2016 | F | N |
| Tg2576 | 159 | + | 129S6 | 11.2 | 08/02/2016 | M | Y |
| Tg2576 | 160 | + | 129S6 | 11.2 | 08/02/2016 | M | Y |
| Tg2576 | 161 | - | 129S6/FVB | 12.2 | 04/05/2015 | F | N |
| Tg2576 | 162 | + | 129S6/FVB | 12.2 | 04/05/2015 | F | Y |
| Tg2576 | 163 | + | 129S6/FVB | 12.2 | 04/05/2015 | F | Y |
| Tg2576 | 164 | - | 129S6/FVB | 12.4 | 03/30/2015 | F | N |
| Tg2576 | 165 | + | 129S6/FVB | 12.6 | 03/24/2015 | F | N |
| Tg2576 | 166 | + | 129S6/FVB | 12.6 | 03/25/2015 | F | N |
| Tg2576 | 167 | + | 129S6/FVB | 12.6 | 03/25/2015 | F | Y |
| Tg2576 | 168 | + | 129S6/FVB | 12.6 | 03/25/2015 | F | Y |
| Tg2576 | 169 | + | 129S6/FVB | 13.2 | 03/27/2015 | F | N |
| Tg2576 | 170 | + | 129S6/FVB | 13.2 | 03/27/2015 | F | N |
| Tg2576 | 171 | - | 129S6/FVB | 13.3 | 03/25/2015 | F | N |
| Tg2576 | 172 | + | 129S6/FVB | 13.3 | 03/25/2015 | F | N |
| Tg2576 | 173 | + | 129S6/FVB | 13.3 | 03/25/2015 | F | Y |
| Tg2576 | 174 | + | 129S6/FVB | 13.3 | 03/25/2015 | F | N |
| Tg2576 | 175 | + | 129S6/FVB | 13.3 | 03/25/2015 | F | N |

| Line | ID | Tg[a] | Background | Age (Mo.)[b] | DOB[c] | Gender[d] | ~56-kDa[e] |
|---|---|---|---|---|---|---|---|
| Tg2576 | 176 | + | 129S6 | 14.7 | 12/28/2002 | M | Y |
| Tg2576 | 177 | + | 129S6 | 14.9 | 12/28/2002 | F | Y |
| Tg2576 | 178 | + | 129S6 | 15.2 | 12/03/2002 | F | Y |
| Tg2576 | 179 | + | 129S6 | 15.2 | 12/03/2002 | F | Y |
| Tg2576 | 180 | + | 129S6 | 16.0 | 02/17/2010 | M | Y |
| Tg2576 | 181 | + | 129S6 | 16.1 | 02/15/2010 | M | Y |
| Tg2576 | 182 | + | B6S | 16.7 | 09/26/2007 | M | Y |
| Tg2576 | 183 | + | 129S6 | 16.9 | 05/11/2011 | M | Y |
| Tg2576 | 184 | + | 129S6 | 17.0 | 05/09/2011 | M | Y |
| Tg2576 | 185 | + | 129S6 | 17.0 | 05/11/2011 | M | N |
| Tg2576 | 186 | + | 129S6 | 17.1 | 05/09/2011 | M | Y |
| Tg2576 | 187 | + | 129S6 | 17.1 | 05/09/2011 | F | Y |
| Tg2576 | 188 | - | 129S6 | 17.1 | 05/09/2011 | M | N |
| Tg2576 | 189 | - | 129S6 | 17.1 | 05/09/2011 | F | N |
| Tg2576 | 190 | - | 129S6 | 17.1 | 05/09/2011 | F | N |
| Tg2576 | 191 | - | 129S6 | 17.1 | 05/09/2011 | M | N |
| Tg2576 | 192 | - | B6S | 17.3 | 09/13/2005 | F | N |
| Tg2576 | 193 | - | B6S | 20.0 | 05/31/2005 | F | N |
| Tg2576 | 194 | + | 129S6 | 21.7 | 02/15/2010 | F | Y |
| Tg2576 | 195 | + | 129S6 | 21.7 | 02/15/2010 | M | Y |
| Tg2576 | 196 | + | 129S6 | 21.7 | 02/16/2010 | F | Y |
| Tg2576 | 197 | + | 129S6 | 21.7 | 02/16/2010 | M | Y |
| Tg2576 | 198 | + | 129S6 | 21.7 | 02/17/2010 | F | Y |
| Tg2576 | 199 | + | 129S6 | 21.7 | 02/17/2010 | M | Y |
| Tg2576 | 200 | - | 129S6 | 21.7 | 02/17/2010 | M | N |
| Tg2576 | 201 | + | 129S6/FVB | 24.9 | 04/23/2008 | M | Y |
| Tg2576 | 202 | + | 129S6/FVB | 24.9 | 04/23/2008 | M | Y |
| Tg2576 | 203 | + | 129S6/FVB | 25.0 | 04/22/2008 | F | Y |
| Tg2576 | 204 | + | 129S6/FVB | 25.0 | 04/22/2008 | F | Y |
| Tg2576 | 205 | + | 129S6/FVB | 25.0 | 04/21/2008 | F | Y |
| Tg2576 | 206 | + | 129S6/FVB | 25.0 | 04/22/2008 | F | Y |
| Tg2576 | 207 | - | 129S6/FVB | 25.0 | 04/22/2008 | F | N |
| Tg2576 | 208 | + | 129S6/FVB | 25.0 | 04/22/2008 | F | Y |
| Tg2576 | 209 | + | 129S6/FVB | 25.0 | 04/22/2008 | M | Y |
| rTg9191 | 210 | + | FVB/129S6 | 24.1 | 10/17/2008 | M | N |
| rTg9191 | 211 | + | FVB/129S6 | 24.1 | 10/17/2008 | M | N |
| rTg9191 | 212 | - | FVB/129S6 | 24.1 | 10/17/2008 | F | N |
| rTg9191 | 213 | - | FVB/129S6 | 24.1 | 10/19/2008 | M | N |
| rTg9191 | 214 | + | FVB/129S6 | 24.1 | 10/19/2008 | F | N |
| rTg9191 | 215 | + | FVB/129S6 | 24.1 | 10/19/2008 | F | N |
| rTg9191 | 216 | + | FVB/129S6 | 24.0 | 10/20/2008 | F | N |
| rTg9191 | 217 | + | FVB/129S6 | 24.0 | 10/20/2008 | M | N |

[a]+, mice that express human amyloid precursor protein; -, mice that do not express human amyloid precursor protein. [b]Mo., months. [c]DOB, date of birth (MM/DD/YYYY). [d]M, male; F, female. [e]Y,

the ~56-kDa, Aβ-containing entities are detected; N, the ~56-kDa, Aβ-containing entities are not detected.

**Table S2 Antibodies used in this study**

| Antibody | Host/Isotype[a] | Epitope[b] | Source |
|---|---|---|---|
| D8Q7I | Rb, IgG | C-terminus of Aβ(x-40) | Cell Signaling Technology, Cat# 12990S; RRID: AB_2798082 |
| Biotinylated D8Q7I | Rb, IgG | C-terminus of Aβ(x-40) | Home prepared from antibody RRID: AB_2798082 |
| D3E10 | Rb, IgG | C-terminus of Aβ(x-42) | Cell Signaling Technology, Cat# 12843S; RRID: AB_2798041 |
| Biotinylated 6E10 | Ms, $IgG_{1,\kappa}$ | Aβ(3-8) | BioLegend, Cat# 803009; RRID: AB_2564656 |
| 82E1 | Ms, $IgG_1$ | N-terminus of Aβ(1-x)/CTFβ | IBL America, Cat# 10323; RRID: AB_10707424 |
| Biotinylated 82E1 | Ms, $IgG_1$ | N-terminus of Aβ(1-x)/CTFβ | IBL America, Cat# 10326; RRID: AB_1540459 |
| HJ2 | Ms, $IgG_{\kappa}$ | C-terminus of Aβ(x-40) | A kind gift from Dr. Dominic Walsh |
| HJ7.4 | Ms, $IgG_{2b,\kappa}$ | C-terminus of Aβ(x-42) | A kind gift from Dr. Dominic Walsh |
| A11 | Rb, IgG | Amyloid oligomers | Rockland, Cat# 200-401-E88; RRID: AB_2612166 |
| A11 | Rb, IgG | Amyloid oligomers | StressMarq Biosciences, Cat# SPC-506D; RRID: AB_10962958 |
| Biotinylated A11 | Rb, IgG | Amyloid oligomers | StressMarq Biosciences, Cat# SPC -506D-BI; RRID: AB_10962958 |
| A11 | Rb, IgG | Amyloid oligomers | Invitrogen, Cat#AHB0052; RRID: AB_2536236 |
| A11 | Rb, IgG | Amyloid oligomers | MilliporeSigma, Cat# AB9234; RRID: AB_570955 |
| Biotin-SP (long spacer) AffiniPure donkey-anti-rabbit IgG (H+L) | Dk, IgG | Heavy and light chains of rabbit IgG | Jackson ImmunoResearch Laboratories, Cat# 711-065-152; RRID: AB_2340593 |

[a]Ms = mouse, Rb = rabbit, Dk = donkey, IgG = immunoglobulin G. [b]Aβ(x-40), human amyloid-β amino acids x-40; Aβ(x-42), human amyloid-β amino acids x-42; Aβ(3-8), human amyloid-β amino acids 3-8; Aβ(1-x), human amyloid-β amino acids 1-x; CTFβ, C-terminal fragment of β-secretase-cleaved human amyloid precursor protein.

**Table S3 Mice and reagents used in immunoprecipitation, immunoaffinity purification, and western blotting**

| Figure | Animal ID | Brain Extracts | Capturing Reagent | Matrix[b] | Detecting Reagent |
|---|---|---|---|---|---|
| 1B, lane 1 “RbIgG/+°” | 99, 125, 126, 127, 132, 133, 134 & 173 | 50 µg each, pooled | RbIgG (3.1µg) | SephG (60 µL) | Biotinylated 6E10 (1:2,500; 400 ng/mL[c]) / Neutravidin-HRP (1:5,000) |
| 1B, lane 3 | 173 | 400 µg | D8Q7I (3.1µg) | SephG (60 µL) | |
| 1B, lane 4 | 99 | 400 µg | D8Q7I (3.1µg) | SephG (60 µL) | |
| 1B, lane 5 | 132 | 400 µg | D8Q7I (3.1µg) | SephG (60 µL) | |
| 1B, lane 6 | 133 | 400 µg | D8Q7I (3.1µg) | SephG (60 µL) | |
| 1B, lane 7 | 134 | 400 µg | D8Q7I (3.1µg) | SephG (60 µL) | |
| 1B, lane 8 | 125 | 400 µg | D8Q7I (3.1µg) | SephG (60 µL) | |
| 1B, lane 9 | 126 | 400 µg | D8Q7I (3.1µg) | SephG (60 µL) | |
| 1B, lane 10 | 127 | 400 µg | D8Q7I (3.1µg) | SephG (60 µL) | |
| 1B, lane 11 | 128 | 400 µg | D8Q7I (3.1µg) | SephG (60 µL) | |
| 1B, lane 12 | 211 | 400 µg | D8Q7I (3.1µg) | SephG (60 µL) | |
| 2B, lane 1 “RbIgG/+°” | 48, 99, 120, 125, 133, 162 & 167 | 51 µg each, pooled | RbIgG (3.1µg) | SephG (60 µL) | Biotinylated 82E1 (1:1,000; 100 ng/mL[c]) / Neutravidin-HRP (1:5,000) |
| 2B, lane 3 | 145 | 360 µg | D8Q7I (3.1µg) | SephG (60 µL) | |
| 2B, lane 4 | 48 | 360 µg | D8Q7I (3.1µg) | SephG (60 µL) | |
| 2B, lane 5 | 167 | 360 µg | D8Q7I (3.1µg) | SephG (60 µL) | |
| 2B, lane 6 | 162 | 360 µg | D8Q7I (3.1µg) | SephG (60 µL) | |
| 2B, lane 7 | 120 | 360 µg | D8Q7I (3.1µg) | SephG (60 µL) | |
| 2B, lane 8 | 99 | 360 µg | D8Q7I (3.1µg) | SephG (60 µL) | |
| 2B, lane 9 | 133 | 360 µg | D8Q7I (3.1µg) | SephG (60 µL) | |
| 2B, lane 10 | 125 | 360 µg | D8Q7I (3.1µg) | SephG (60 µL) | |
| 2B, lane 11 | 128 | 360 µg | D8Q7I (3.1µg) | SephG (60 µL) | |
| 3A | 80, 81, 85, 91 & 106 | 100 µg each, pooled[a] | N/A | N/A | Biotinylated 82E1 (1:1,000; 100 ng/mL[c]) / Neutravidin-HRP (1:5,000) |
| 3C | 80, 81, 85, 91 & 106 | 100 µg each, pooled[a] | N/A | N/A | Biotinylated D8Q7I (1:10,000; 19 ng/mL[c]) / Neutravidin-HRP (1:5,000) |

| Figure | Animal ID | Brain Extracts | Capturing Reagent | Matrix[b] | Detecting Reagent |
| --- | --- | --- | --- | --- | --- |
| 4, lane "TBS/-" | 157 | 375 µg | D8Q7I (3.1 µg) | SephG (80 µL) | Biotinylated 82E1 (1:1,000; 100 ng/mL[c]) / Neutravidin-HRP (1:5,000) |
| 4, lane "TBS/+" | 48, 99, 125 & 133 | 94 µg each, pooled | D8Q7I (3.1 µg) | SephG (80 µL) | |
| 4, lane "urea/+" | 48, 99, 125 & 133 | 94 µg each, pooled | D8Q7I (3.1 µg) | SephG (80 µL) | |
| 4, lane "GuHCl/+" | 48, 99, 125 & 133 | 94 µg each, pooled | D8Q7I (3.1 µg) | SephG (80 µL) | |
| 4, lane "HFIP/+" | 48, 99, 125 & 133 | 94 µg each, pooled | D8Q7I (3.1 µg) | SephG (80 µL) | |
| 5A, lane "+" | 176, 178, 179, 180, 196, 197, 198 & 199 | 62.5 µg each, pooled | D8Q7I (7.8 µg) | DynaG (125 µL) | Biotinylated 82E1 (1:1,000; 100 ng/mL[c]) / Neutravidin-HRP (1:5,000) |
| 5A, lane "-" | 186, 187, 188, 189 & 200 | 100 µg each, pooled | D8Q7I (7.8 µg) | DynaG (125 µL) | |
| 5B, lane "+" | 176, 178, 179, 180, 196, 197, 198 & 199 | 62.5 µg each, pooled | D8Q7I (7.8 µg) | DynaG (125 µL) | Biotinylated A11 (1:1,000) / Neutravidin-HRP (1:5,000) |
| 5B, lane "-" | 186, 187, 188, 189 & 200 | 100 µg each, pooled | D8Q7I (7.8 µg) | DynaG (125 µL) | |
| 5C, lane "+" | 176, 178, 179, 180, 196, 197, 198 & 199 | 125 µg each, pooled | D8Q7I (31 µg) | DynaG (500 µL) | |
| 5C, lane "-" | 186, 187, 188, 189 & 200 | 200 µg each, pooled | D8Q7I (31 µg) | DynaG (500 µL) | |
| S1, lane "14-16, + (left)" | 177 | 400 µg | D8Q7I (3.1µg) | SephG (50 µL) | Biotinylated 82E1 (1:1,000; 100 ng/mL[c]) / Neutravidin-HRP (1:5,000) |
| S1, lane "14-16, + (right)" | 181 | 400 µg | D8Q7I (3.1µg) | SephG (50 µL) | |
| S1, lane "21, -" | 200 | 400 µg | D8Q7I (3.1µg) | SephG (50 µL) | |
| S1, lane "21, + (left)" | 194 | 400 µg | D8Q7I (3.1µg) | SephG (50 µL) | |
| S1, lane "21, + (right)" | 195 | 400 µg | D8Q7I (3.1µg) | SephG (50 µL) | |

| Figure | Animal ID | Brain Extracts | Capturing Reagent | Matrix[b] | Detecting Reagent |
|---|---|---|---|---|---|
| S2A-B, lane "-" | 128, 145 & 157 | 127 µg each, pooled | 82E1 (3 µg) | SephG (60 µL) | D8Q7I (1:2,500; 245 ng/mL[c]) / biotinylated donkey-anti-rabbit IgG (1:50,000) / Neutravidin-HRP (1:5,000) |
| S2A-B, lane "+ (left)" | 133, 134, 153, 155, 162 & 167 | 63 µg each, pooled | 82E1 (3 µg) | SephG (60 µL) | |
| S2A-B, lane "+ (right)" | 48, 99, 120, 125, 126 & 127 | 63 µg each, pooled | 82E1 (3 µg) | SephG (60 µL) | |
| S2C-D, lane "14-15, -" | 200 | 400 µg | 82E1 (3 µg) | SephG (50 µL) | Biotinylated D8Q7I (1:10,000; 19 ng/mL[c]) / Neutravidin-HRP (1:5,000) |
| S2C-D, lane "14-15, + (left)" | 179 | 400 µg | 82E1 (3 µg) | SephG (50 µL) | |
| S2C-D, lane "14-15, + (right)" | 176 | 400 µg | 82E1 (3 µg) | SephG (50 µL) | |
| S2C-D, lane "25, -" | 207 | 400 µg | 82E1 (3 µg) | SephG (50 µL) | |
| S2C-D, lane "25, + (left)" | 203 | 400 µg | 82E1 (3 µg) | SephG (50 µL) | |
| S2C-D, lane "25, + (right)" | 205 | 400 µg | 82E1 (3 µg) | SephG (50 µL) | |
| S3A, lane 1 "RbIgG/+°" | 48, 99, 120, 125, 133, 162 & 167 | 51 µg each, pooled | D3E10 (168 ng) | SephG (60 µL) | Biotinylated 6E10 (1:2,500; 400 ng/mL[c]) / Neutravidin-HRP (1:5,000) |
| S3A, lane 3 | 145 | 360 µg | D3E10 (168 ng) | SephG (60 µL) | |
| S3A, lane 4 | 48 | 360 µg | D3E10 (168 ng) | SephG (60 µL) | |
| S3A, lane 5 | 167 | 360 µg | D3E10 (168 ng) | SephG (60 µL) | |
| S3A, lane 6 | 162 | 360 µg | D3E10 (168 ng) | SephG (60 µL) | |
| S3A, lane 7 | 120 | 360 µg | D3E10 (168 ng) | SephG (60 µL) | |
| S3A, lane 8 | 99 | 360 µg | D3E10 (168 ng) | SephG (60 µL) | |
| S3A, lane 9 | 133 | 360 µg | D3E10 (168 ng) | SephG (60 µL) | |
| S3A, lane 10 | 125 | 360 µg | D3E10 (168 ng) | SephG (60 µL) | |
| S3A, lane 11 | 128 | 360 µg | D3E10 (168 ng) | SephG (60 µL) | |

| Figure | Animal ID | Brain Extracts | Capturing Reagent | Matrix[b] | Detecting Reagent |
|---|---|---|---|---|---|
| S3B-C, lane "-" | 128, 145 & 157 | 127 µg each, pooled | 82E1 (3 µg) | SephG (60 µL) | D3E10 (1:2,500; 13.4 ng/mL[c]) / biotinylated donkey-anti-rabbit IgG (1:50,000) / Neutravidin-HRP (1:5,000) |
| S3B-C, lane "+ (left)" | 48, 99, 120, 125, 126 & 127 | 63 µg each, pooled | 82E1 (3 µg) | SephG (60 µL) | |
| S3B-C, lane "+ (right)" | 133, 134, 153, 155, 162 & 167 | 63 µg each, pooled | 82E1 (3 µg) | SephG (60 µL) | |
| S4, lane "HJ2, +" | 173 | 200 µg | HJ2 (5 µg) | DynaG (50 µL) | Biotinylated 6E10 (1:2,500; 400 ng/mL[c]) / Neutravidin-HRP (1:5,000) |
| S4, lane "HJ2, -" | 171 | 200 µg | HJ2 (5 µg) | DynaG (50 µL) | |
| S4, lane "HJ2, +" | 173 | 200 µg | HJ7.4 (5.3 µg) | DynaG (50 µL) | |
| S4, lane "HJ2, -" | 171 | 200 µg | HJ7.4 (5.3 µg) | DynaG (50 µL) | |
| S4, lane "HJ2, +" | 173 | 200 µg | D8Q7I (3.1 µg) | DynaG (50 µL) | |
| S4, lane "HJ2, -" | 171 | 200 µg | D8Q7I (3.1 µg) | DynaG (50 µL) | |
| S5 | 80, 81, 85, 91 & 106 | 100 µg each, pooled[a] | N/A | N/A | Neutravidin-HRP (1:5,000) |
| S6, lanes "+" | 176, 178, 179, 180, 196, 197, 198 & 199 | 62.5 µg each, pooled | D8Q7I (7.8 µg) | DynaG (125 µL) | Neutravidin-HRP (1:5,000) |
| S6, lanes "-" | 186, 187, 188, 189 & 200 | 100 µg each, pooled | D8Q7I (7.8 µg) | DynaG (125 µL) | |
| S7A-D, lanes "-" | 186, 187, 188, 189 & 200 | 10 µg each, pooled | N/A | N/A | A11 (1:5,000) / biotinylated donkey-anti-rabbit IgG (1:50,000) / Neutravidin-HRP (1:5,000) |
| S7A-D, lanes "+" | 176, 178, 179, 180, 196, 197, 198 & 199 | 6.25 µg each, pooled | N/A | N/A | |

| Figure | Animal ID | Brain Extracts | Capturing Reagent | Matrix[b] | Detecting Reagent |
|---|---|---|---|---|---|
| S8, lane 3 "RbIgG/+°" | 182, 183, 184, 185, 190 & 191 | 67 µg each, pooled | RbIgG (3.1µg) | SephG (50 µL) | Biotinylated 82E1 (1:1,000; 100 ng/mL[c]) / Neutravidin-HRP (1:5,000) |
| S8, lane 4 "-" | 189 | 400 µg | D8Q7I (3.1 µg) | SephG (50 µL) | |
| S8, lane 5 "+" | 182 | 400 µg | D8Q7I (3.1 µg) | SephG (50 µL) | |
| S8, lane 6 "+" | 183 | 400 µg | D8Q7I (3.1 µg) | SephG (50 µL) | |
| S8, lane 7 "+" | 184 | 400 µg | D8Q7I (3.1 µg) | SephG (50 µL) | |
| S8, lane 8 "+" | 185 | 400 µg | D8Q7I (3.1 µg) | SephG (50 µL) | |
| S8, lane 9 "+" | 190 | 400 µg | D8Q7I (3.1 µg) | SephG (50 µL) | |
| S8, lane 10 "+" | 191 | 400 µg | D8Q7I (3.1 µg) | SephG (50 µL) | |

[a]The western blotting (WB) analyses of fractions of size-exclusion chromatography was obtained from fractionating the pooled material, 500 µg in total; the WB analyses of input was obtained from 50 µg of pooled material (10 µg from each animal). [b]The volumes listed in this column are the volumes of beads in slurry. The net volumes of beads are approximately 50% (SephG) or 10% (DynaG) of the reported volumes. SephG, Protein G Sepharose 4 Fast Flow resin; DynaG, Dynabeads Protein G. [c]The final concentration of the primary antibody is shown.
